# Unlocking biological insight from single-cell data with an interpretable dual-stream foundation model

**DOI:** 10.1101/2025.09.05.674596

**Authors:** Honglie Guo, Qinghang Cui, Xiang Zhang, Chaowei Chen, Weihua Zheng, Changfeng Cai, Xinyi Wang, Shunfang Wang

## Abstract

Deep learning foundation models are revolutionizing single-cell biology, yet learning holistic and discriminative representations from complex, high-dimensional data remains a central challenge. Although Transformer-based single-cell language models have shown significant progress, they typically rely on a single input-encoding scheme, a practice that results in the loss of critical gene expression information and hinders the effective learning of global cellular representations. To address these challenges, we introduce scDMC, an innovative single-cell Dual-stream Masked Contrastive pre-training framework designed to synergistically optimize information fidelity at both the gene and cellular levels. Pre-trained on only 2 million cells far fewer than the datasets used by mainstream models, scDMC sets a new state-of-the-art in multiple benchmark tasks, including cell annotation, clustering, and data integration. More importantly, we demonstrate that scDMC can uncover functional gene modules, infer cell-type-specific regulatory networks in a data-driven manner, and exhibits a high degree of biological interpretability.

## 1 Introduction

The rapid advancement of single-cell RNA sequencing (scRNA-seq) technology allows us to resolve transcriptomes at an unprecedented resolution, profoundly illuminating the cellular heterogeneity and dynamic changes inherent in complex biological processes [1–4]. However, the analysis of single-cell data remains hampered by inherent noise, platform-specific technical variations, and the characteristic high-dimensionality and sparsity of the data. These challenges hinder accurate downstream analyses such as cell-type annotation [5–7] and make it difficult to correct for batch effects arising from different sequencing technologies [8–11].

Inspired by successes in natural language processing, Transformer-based foundation models have emerged as a leading strategy to tackle these challenges [12, 13]. Following the tremendous success of pre-trained language models (PLMs) [14] like BERT [15] and the GPT series [16–19], the “pre-train, fine-tune” paradigm has been actively adopted in the field of single-cell data analysis [20]. By pre-training on large-scale, unlabeled data, models such as scGPT [21], Geneformer [22], and scFoundation [23] have demonstrated great potential across numerous tasks. These single-cell language models (scLMs) [24] analogize cells to “sentences” and genes to “words” attempting to learn the “grammatical rules” of life. Other work, such as LangCell [25], has even explored unifying single-cell and natural language representations to enhance model comprehension with external knowledge.

Most of these models adopt masked language modeling (MLM) from BERT [15] as their primary pre-training objective. However, the standard Transformer architecture faces a bottleneck of quadratic computational and memory complexity when processing the ultra-long sequences typical of single-cell transcriptomics, which can include tens of thousands of genes [26, 27]. Consequently, researchers have actively explored more efficient Transformer variants and attention mechanisms. For instance, scBERT [28] employed the Performer [29] architecture to reduce attention complexity, while scGPT [21] leveraged the highly efficient Flash Attention [30] for model training. Concurrently, other efforts have begun to focus on newer state-space models (SSMs), such as the Mamba [31] architecture used in GeneMamba [32]. Geneformer-95M [33] has also explored resource-efficient model fine-tuning for downstream tasks, quantizing the model to 4-bit precision using low-rank adapters (QLoRA) to reduce computational demands.

Despite this initial success, a fundamental dilemma persists for existing scLMs: how to encode non-sequential single-cell expression profiles into a sequence that a model can understand without substantial information loss. Current approaches present a difficult trade-off. Rank-based methods [22], for instance, discretize the transcriptome by sorting genes according to their expression levels and treating the resulting sequence of gene IDs as input tokens; while this effectively captures the relative functional context, it fundamentally discards all information about the absolute magnitude of expression. Conversely, expression binning [21, 28] attempts to retain magnitude by grouping values into discrete bins and using these as positional information, but this comes at the cost of reduced data resolution and a performance highly dependent on the chosen binning strategy. While value projection methods [34] use a multilayer perceptron on continuous expression values to avoid this loss of resolution, their universal effectiveness across diverse biological scenarios has yet to be fully established. Furthermore, most models rely excessively on the singular MLM pre-training task and lack an explicit mechanism for learning the higher-order features that distinguish global cell states.

To address both of these core bottlenecks simultaneously, we developed scDMC (a single-cell Dual-stream MLM and Contrastive learning framework). At the heart of scDMC is an innovative dual-stream pre-training framework that, for the first time, preserves both the relative importance and absolute magnitude of gene expression by processing two complementary gene sequences in parallel. Furthermore, we introduce momentum contrastive learning [35, 36], which ingeniously leverages the two cell-level views generated by the dual streams as a natural positive pair, thereby driving the model to learn a more discriminative global cell representation space. This design, which integrates local gene context with global cell representation, not only enables scDMC to excel in tasks like cell-type annotation and batch integration but also endows it with the ability to conduct biological exploration at a higher resolution. Additionally, we have incorporated efficient training strategies [37], such as FlashAttention [30] and ALiBi [38] positional encoding, to ensure both efficiency and performance when processing long sequences.

In this work, we systematically demonstrate the powerful performance and remarkable efficiency of the scDMC framework. We show that scDMC achieves state-of-the-art performance across a range of challenging downstream tasks, including cell-type annotation, unsupervised clustering, and multi-batch data integration. Remarkably, these results were achieved by pre-training on just 2 million cells (a dataset size substantially smaller than those used by comparable models) demonstrating the framework’s superior data efficiency and design advantages. Finally, we delve into the biological meaning of the embedding space learned by scDMC, demonstrating that it not only uncovers functional gene modules and infers cell-type-specific regulatory networks but also that its decision-making process is highly interpretable, offering a powerful new paradigm for data-driven biological discovery in single-cell data.

## 2 Results

### 2.1 The scDMC pre-training framework

The core challenge in single-cell language modeling stems from the dual nature of cellular information: a model must be faithful to the complex contextual relationships among local genes while simultaneously capturing the unique signature of the global cell identity. Existing models often struggle to reconcile these two aspects. To address this challenge, we developed scDMC, an innovative pre-training framework designed to learn comprehensive and robust biological representations at both the gene and cell levels (Fig. 1). This framework’s design synergistically addresses the field’s core challenges through three key innovations.

**Fig. 1.**
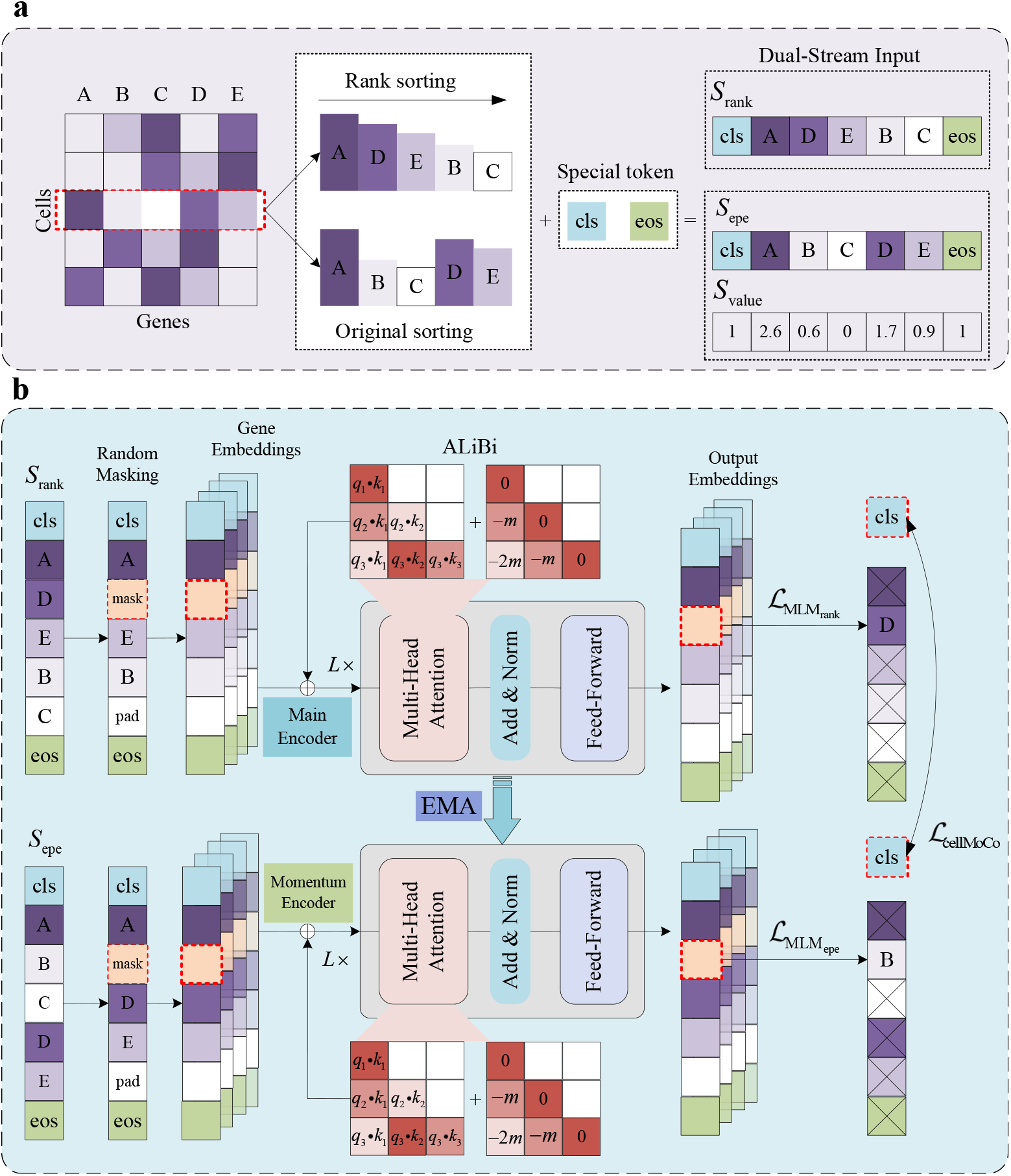
The scDMC model architecture. **a**, Dual-stream input construction. For a single cell, two input sequences are generated: **S**_rank_, which is based on gene expression ranking, and **S**_epe_, **S**_value_, which preserves the original gene order and carries expression value information. **b**, The model pretraining process. These two sequences are masked and then fed into the main encoder. Concurrently, the momentum encoder also processes both input streams to generate target representations. The model’s total loss is composed of two components: the dual-stream MLM loss, where the loss for the second stream is weighted by expression values, and the contrastive loss, calculated by contrasting the **[CLS]** representations output by the main and momentum encoders.

First, at the gene level, we design a unique dual-stream masked language modeling (Dual-Stream MLM) strategy to overcome the information loss inherent in single-encoding schemes. One data stream processes gene sequences ranked by expression level, enabling it to focus on the relative importance and functional context of genes within a specific cellular state. Concurrently, a second processing stream maintains the original gene order while employing an expression-weighted loss function to preserve absolute expression values, critical biological information that is typically lost in rank-based approaches. Through joint optimization of these complementary representations within a unified framework, scDMC produces more comprehensive and biologically interpretable gene embeddings than conventional single-stream methods.

Second, to enable the model with the ability to learn highly discriminative cell-level global representations, we introduced a cell momentum contrastive learning mechanism. Here, we innovatively leverage the unique advantage of the dual-stream architecture: the two embedding representations generated from the same cell across the different data streams are treated as a natural, high-quality “positive pair”. This self-supervised task drives the model to learn a structured embedding space where different views of the same cell are pulled closer, while views of different cells are pushed apart. This critically enhances the model’s ability to distinguish subtle differences between cellular states and types, a key factor in its superior performance on downstream tasks.

Finally, to ensure the computational feasibility and scalability of this complex framework when processing long sequences containing tens of thousands of genes, we integrated several key architectural optimizations [39]. Critically, to accommodate the non-sequential and variable-length nature intrinsic to single-cell data, we replaced traditional positional encodings with Attention with Linear Biases (ALiBi). ALiBi does not rely on fixed positional embeddings; instead, it dynamically adjusts attention scores based on the relative distance between tokens, making it more flexible and effective at capturing the complex relative importance and functional associations among genes.

The dataset used for scDMC pre-training comprises 2 million single-cell transcriptomes from healthy human blood, sourced from the CELLxGENE Discover Census database [40]. To maximally preserve the original biological signals and avoid filtering-induced biases, we adopted a minimalist pre-processing strategy. No upstream cell quality control [41] (e.g., filtering by mitochondrial gene percentage) or feature selection [42] (e.g., identifying highly variable genes) was performed when constructing the pre-training corpus. This design forces the model to learn intrinsic biological principles directly from near-raw data. The cell type composition and heterogeneity of the final pre-training dataset are detailed in Supplementary Fig. 4 and Supplementary Fig. 5. Following pre-training, we applied the scDMC model to multiple downstream tasks.

To evaluate the generalization capabilities and robustness of scDMC, we selected six public benchmark datasets with high-quality cell type annotations for testing: Zheng68K [43], PBMC10k [44], Immune Human [45], BMMC [46], MS [47], and Myeloid [48]. These datasets were chosen to cover a diverse range of biological contexts, including different tissue sources, sequencing platforms, and health statuses (Supplementary Table 2).

### 2.2 scDMC achieves superior performance in cell type annotation

Accurate cell type identification serves as the gold standard for validating the effectiveness of single-cell representation models and is one of the most central tasks for their evaluation. To fine-tune the model for this task, we initialized it with the pretrained scDMC weights and added a final linear layer for classification. During this process, the model takes the ranked single-cell gene expression sequence as input and learns to predict specific cell types (e.g., B cells, T cells).

To rigorously evaluate scDMC’s performance, we selected four benchmark datasets characterized by high diversity in tissue source, sequencing technology, and health status. Across all tests, scDMC consistently achieved state-of-the-art (SOTA) or highly competitive performance, demonstrating particularly strong results in the Macro F1-Score, a metric that provides an unbiased measure of a model’s performance on imbalanced datasets, typical of real-world biological data, by weighting all cell classes equally (Fig. 2a). On the comprehensive BMMC dataset, scDMC achieved a remarkable 5.18% improvement in Macro F1-Score over the next-best model, clearly demonstrating its robustness in handling mixed data from multiple technology platforms and batches (Fig. 2b). Similarly, on the MS (multiple sclerosis) dataset, which includes disease-state samples, scDMC also exhibited top-tier performance (Fig. 2a, 2b). This indicates that the cellular representations learned by scDMC can effectively distinguish the subtle cellular differences between healthy and diseased states, highlighting its significant potential for clinical applications.

**Fig. 2.**
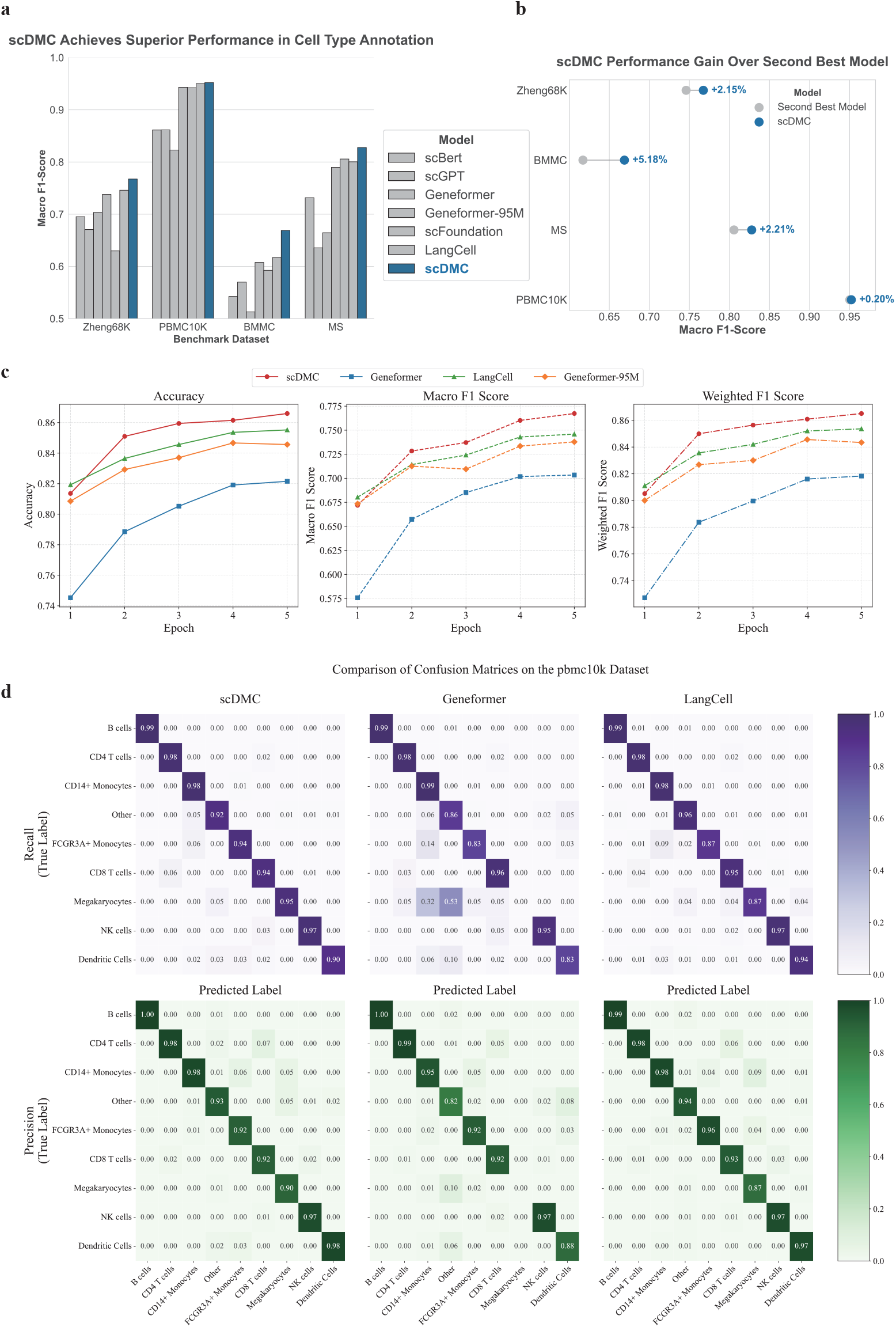
scDMC achieves superior performance in cell type annotation. **a**, Comparison of Macro F1-Scores for scDMC and other leading single-cell language models on the cell type annotation task across four representative benchmark datasets (Zheng68K, PBMC10K, BMMC, MS). scDMC (dark blue bars) achieves the top performance in all datasets, demonstrating its powerful generalization and robustness. **b**, Percentage improvement in Macro F1-Score of scDMC compared to the next-best model for each dataset. The results show that scDMC achieves performance gains of 2.15% (Zheng68K), 0.20% (PBMC10K), 5.18% (BMMC), and 2.21% (MS), proving its significant superiority. **c**, Performance curves of different models with increasing fine-tuning epochs on the highly challenging Zheng68K dataset. From left to right: Accuracy, Macro F1-Score, and Weighted F1-Score. scDMC (red dashed line) exhibits the fastest convergence speed and the highest performance ceiling across all metrics. **d**, Comparison of normalized confusion matrices for scDMC and two representative models (Geneformer, LangCell) on the PBMC10k dataset. The matrices are normalized by true labels (Recall, top row, purple) and predicted labels (Precision, bottom row, green), respectively. The values on the diagonal represent the recall or precision for each class, with color intensity corresponding to the value. The results clearly show that scDMC classifies nearly all cell types perfectly, with off-diagonal regions close to zero, indicating extremely high classification accuracy and low class confusion. In contrast, other models exhibit significant confusion for several classes.

The robustness of scDMC is particularly evident when handling highly challenging datasets. In the Zheng68K dataset, which is notorious for its severe cell-type imbalance and high similarity between subtypes, scDMC achieved a cell type identification accuracy of 86.60% and a Macro F1-Score of 0.767, surpassing the next-best model by 2.15 percentage points (Supplementary Table 3). In both performance and learning efficiency, it comprehensively outperformed existing models (Fig. 2c). Notably, the learning curve for scDMC not only reached a higher performance ceiling but also consistently maintained its lead throughout the fine-tuning process. This suggests that scDMC can learn the key features of cell types more efficiently and stably, demonstrating a superior ability to handle class imbalance and accurately identify rare cell subpopulations (Supplementary Fig. 6).

To delve deeper into the source of scDMC’s performance advantages, we visualized its classification confusion matrix on the PBMC10k dataset (Fig. 2d). The result is strikingly clear: scDMC classified nearly all cell types perfectly, with diagonal values for both the Recall and Precision matrices approaching 1 and virtually no confusion in the off-diagonal regions. In sharp contrast, other models, such as Geneformer, exhibited significant difficulty in distinguishing between similar cell types, particularly showing a notable performance bottleneck in identifying Megakaryocytes.

This result provides compelling evidence that scDMC’s unique dual-stream pretraining framework and momentum contrastive learning mechanism successfully generate highly discriminative cellular representations. These representations can clearly distinguish even closely related biological cell subtypes, thereby resolving the classification ambiguities present in other models. Consequently, scDMC’s ability to provide precise and reliable cell annotation across diverse and complex biological scenarios establishes it as a powerful tool for advancing future biomedical research.

### 2.3 scDMC’s embeddings capture intrinsic cellular and biological structures

Beyond its performance in supervised tasks, a truly powerful foundation model should be able to uncover the intrinsic biological identities within data based on its structure alone, in an unsupervised setting.

To begin, we directly investigated the structure of the native high-dimensional space by performing hierarchical clustering on the original 512-dimensional cell embeddings generated by scDMC for the PBMC10k dataset. The result is striking: even without any dimensionality reduction or non-linear transformation, cells with the same biological identity already form well-defined, highly pure, contiguous blocks (Fig. 3d). This discovery provides direct evidence that the representation space learned by scDMC is, in itself, a structured and biologically meaningful space where the distances between cells directly reflect their biological similarity.

**Fig. 3.**
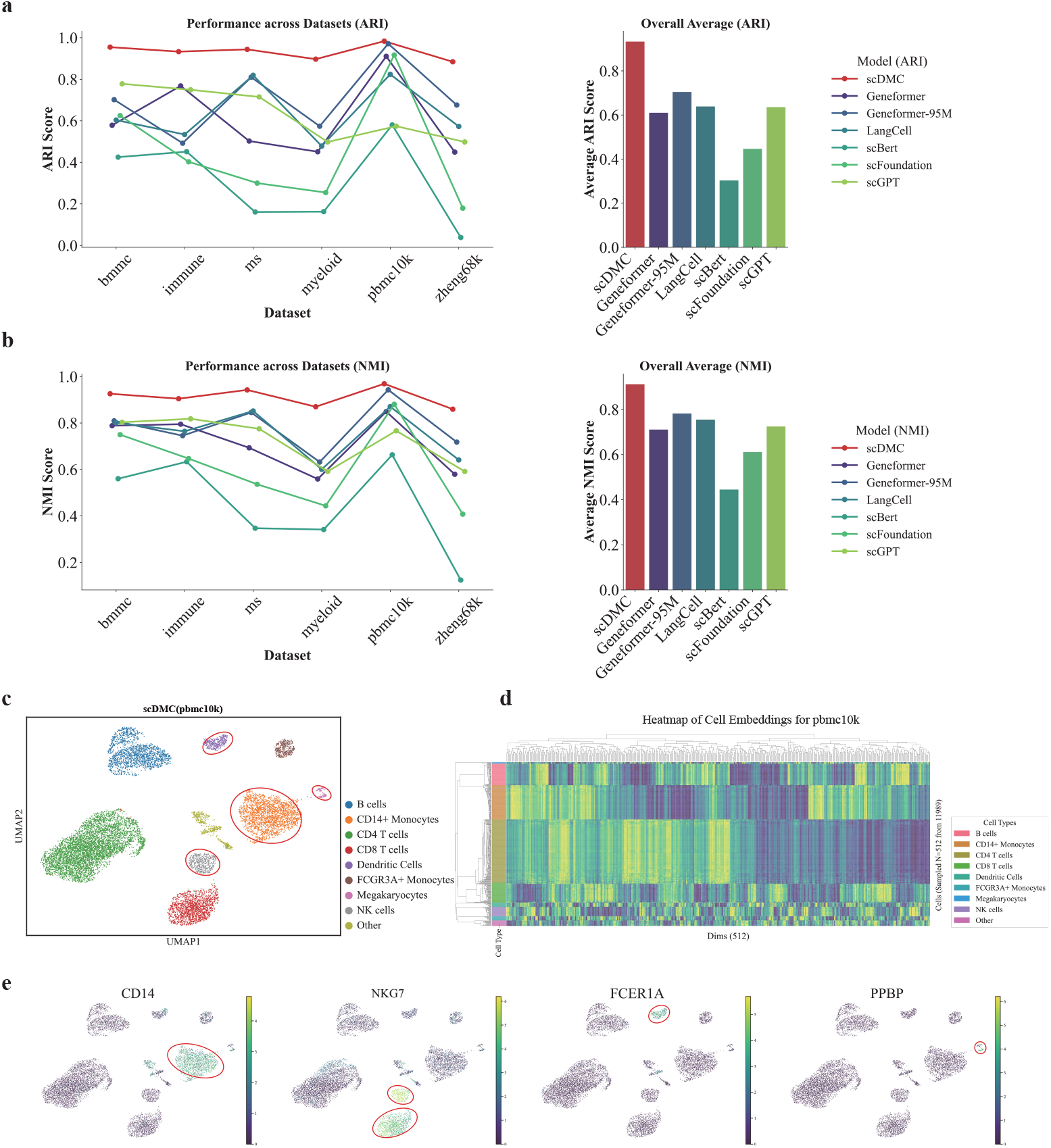
scDMC’s embeddings capture intrinsic cellular and biological structures. **a, b**, Performance of Adjusted Rand Index (ARI) (a) and Normalized Mutual Information (NMI) (b) after unsupervised Leiden clustering on cell embeddings generated by various models across six benchmark datasets. The line plots on the left show performance stability across datasets, while the bar plots on the right represent the average scores over all datasets. scDMC (red) exhibits the most stable and superior performance in all tests, demonstrating that its learned cell representations have the highest consistency with true cell population divisions. **c**, UMAP visualization of cell embeddings for the PBMC10k dataset learned by scDMC. Each point represents a cell, colored by its true cell type label. The results show that major cell lineages form well-defined, distinct clusters, and even rare cell types (such as Dendritic cells and Megakaryocytes, circled in red) are effectively separated. **d**, Heatmap of hierarchical clustering on the raw, un-reduced 512-dimensional scDMC embeddings for 512 cells randomly sampled from the PBMC10k dataset (11,989 cells). Each row represents a cell, colored by the true cell type label on the left; each column represents a dimension of the embedding vector. The plot clearly shows that cells with the same biological identity naturally cluster together, forming pure blocks, which directly demonstrates the highly intrinsic biological structure of the high-dimensional representation space generated by scDMC. **e**, Projection of key marker gene expression levels onto the UMAP plot from (c). Color intensity represents the level of gene expression. The results show that the expression patterns of marker genes are highly consistent with the cell clusters defined by scDMC: CD14 expression defines the CD14+ Monocytes cluster, NKG7 defines the NK cells cluster, FCER1A defines the Dendritic cells cluster, and PPBP defines the Megakaryocytes cluster (relevant clusters are circled in red).

This high-quality intrinsic structure was also fully validated in quantitative unsupervised clustering evaluations. We extracted the cell embedding representations and applied the standard graph-based clustering algorithm, Leiden [49], evaluating the results using the Adjusted Rand Index (ARI) and Normalized Mutual Information (NMI) (Supplementary Information S6.2). The results show that the clustering performance of scDMC (red) comprehensively and consistently surpassed all other models across all tested datasets, with its average ARI and NMI scores ranking highest (Fig. 3a, 3b). It consistently demonstrated superior performance across all test demonstrated superior performance across all test

This advantage becomes even more intuitive when these high-quality embeddings are projected into two-dimensional space for visualization. The UMAP [50] generated from scDMC embeddings not only clearly separates the major cell lineages but, more importantly, it precisely preserves the fine-grained topological structure of each cell type, such as clearly distinguishing different states of T cells and monocytes (Fig. 3c). By projecting the expression of key marker genes onto the UMAP plot, we found that their expression patterns highly correspond with the cell clusters defined by scDMC (Fig. 3e). For instance, CD14 expression is highly concentrated in the CD14+ Monocytes cluster. NKG7, a marker for Natural Killer (NK) cells and activated cytotoxic T cells, shows high expression levels in both the NK cells and CD8 T cells clusters, which is in complete accordance with established biological facts [51]. Furthermore, FCER1A defines the Dendritic cells cluster, while the expression of PPBP, a marker for Megakaryocytes, is precisely confined to the small and distinct Megakaryocytes cluster. This validates the accuracy of the cell representations learned by scDMC from a biological function perspective.

### 2.4 scDMC effectively integrates heterogeneous datasets and removes batch effects

In real-world biomedical research, single-cell data often originate from different donors, technology platforms, or experimental conditions. The resulting technical variations, known as “batch effects”, are a primary obstacle to data integration and the extraction of biological insights [52, 53]. The same absolute value may represent a different biological “semantic meaning” in the context of a different sequencing batch. A powerful foundation model must be able to overcome this technical noise while preserving true biological heterogeneity. Our research demonstrates that scDMC excels at this critical task.

We systematically evaluated scDMC’s integration capabilities by integrating two highly heterogeneous and challenging datasets: the BMMC dataset, comprising 12 different donor batches, and the Immune dataset, comprising 10 different technology platform batches. The qualitative UMAP visualization results are strikingly clear: in the embedding space generated by scDMC, cells from different batches are uniformly mixed, completely eliminating any signs of batch-specific clustering in both the BMMC (Fig. 4a) and Immune (Fig. 4b) datasets. Simultaneously, the biological identity of the cells is perfectly preserved, as evidenced by the high concordance between the unsupervised clustering results and the true cell types. This ideal state of being “mixed by batch, separated by type” intuitively demonstrates that the representations learned by scDMC are highly robust to technical noise.

**Fig. 4.**
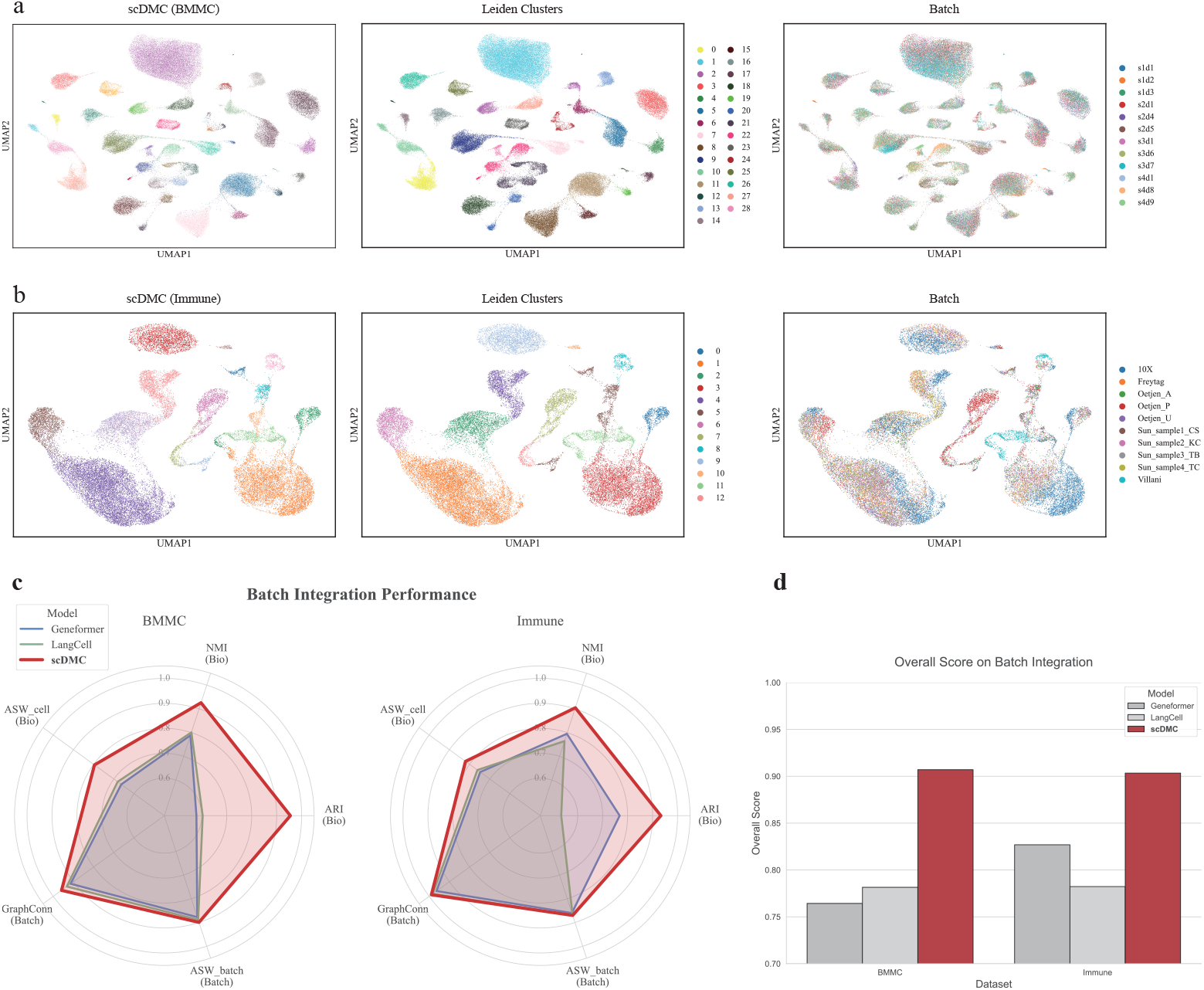
scDMC achieves state-of-the-art data integration and effectively removes batch effects. **a, b**, UMAP visualizations of cell embeddings learned by scDMC on two highly heterogeneous datasets: BMMC (a, containing 12 donor batches) and Immune (b, containing 10 technology platform batches). The left panel is colored by true cell type labels, demonstrating the conservation of biological structure. The middle panel is colored by unsupervised Leiden clustering results, showing high concordance between clusters and true cell types. The right panel is colored by experimental batch, clearly showing that cells from different batches are uniformly mixed with no obvious batch-specific clustering. Together, these panels demonstrate that scDMC achieves the ideal integration effect of being “mixed by batch, separated by type”. **c**, Radar plots showing the quantitative evaluation of data integration performance for different models using the scIB framework on the BMMC (left) and Immune (right) datasets. The plots assess performance across five key metrics: three measuring biological conservation (NMI (Bio), ARI (Bio), ASW cell (Bio)) and two evaluating batch effect correction (ASW batch (Batch), GraphConn (Batch)). The largest area enclosed by scDMC (red) indicates its superior comprehensive performance compared to other models across all evaluation dimensions. **d**, Comparison of the final integration Overall Score for different models on the BMMC and Immune datasets. The results again quantitatively confirm that the integration performance of scDMC (red) is significantly superior to all other models on both datasets.

For rigorous quantitative evaluation, we employed the widely recognized scIB [45] benchmark framework, which comprehensively assesses integration performance from two complementary dimensions: biological information conservation and batch effect correction (Supplementary Information S6.2). The radar plots clearly show that the performance of scDMC (red) comprehensively surpasses that of other models across all key metrics (Fig. 4c). This indicates that scDMC maximally preserves the original biological structure of the cells while effectively removing batch effects. Ultimately, scDMC’s Overall Score on both the BMMC and Immune datasets was significantly higher than all other compared models (Fig. 4d), once again quantitatively confirming its status as a state-of-the-art data integration tool. This powerful integration capability establishes it as an ideal foundation model for constructing large-scale, crossspecies, and cross-disease “single-cell atlases”, paving the way for scientific discovery using vast public data resources.

### 2.5 Uncovering functional gene modules from scDMC embeddings

An outstanding single-cell language model must not only perform accurate cell-level classification but also delve into the gene level to reveal the underlying biological organizational patterns. We demonstrate that scDMC can successfully discover functionally related gene modules and infer cell-type-specific gene regulatory networks in a data-driven manner.

By performing unsupervised clustering on the contextual gene embeddings generated by scDMC, we were able to identify functionally cohesive gene modules within specific cell types (Fig. 5a). Functional enrichment analysis of these modules clearly shows that scDMC can deconstruct complex cellular biological functions into multiple, distinct core programs. For instance, in CD8+ T cells, we identified one module highly associated with adaptive immunity and antigen presentation (Module 4) and another closely linked to ribosome function (Module 3) (Fig. 5c). In hematopoietic stem cells (HSCs), the identified modules corresponded to key processes for maintaining their stemness, such as protein homeostasis and stress response (Fig. 5d). This capacity for data-driven functional deconstruction demonstrates that the gene-level representations learned by scDMC are deeply biologically meaningful.

**Fig. 5.**
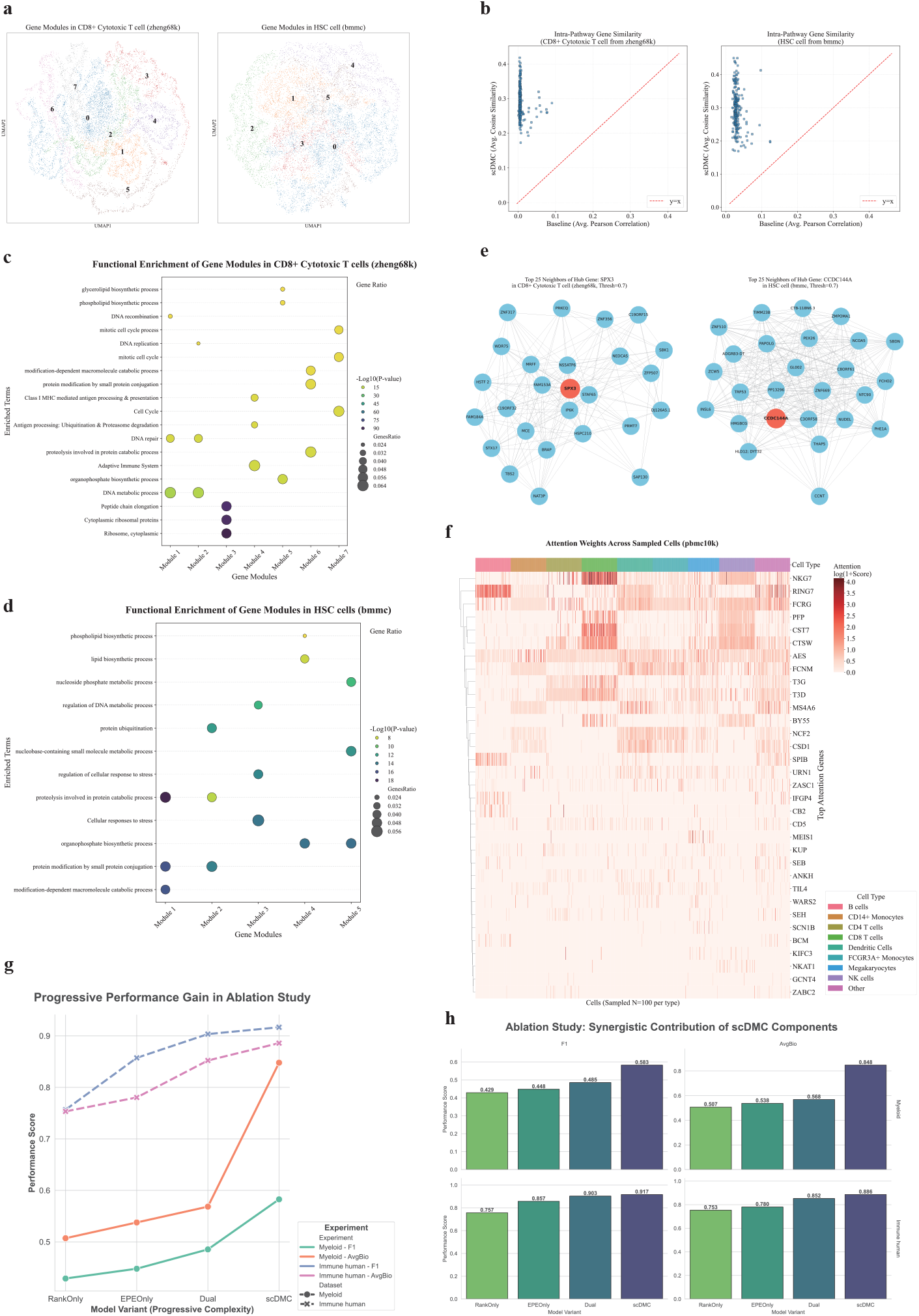
scDMC reveals multi-level biological structures and is interpretable. **a**, Functional gene modules identified by unsupervised clustering on gene embeddings generated by scDMC in CD8+ T cells (left) and hematopoietic stem cells (HSCs) (right). Each point represents a gene, colored by its module assignment. The distinct modular separation indicates that scDMC’s gene embedding space has an intrinsic functional structure. **b**, Quantitative comparison of the internal consistency of scDMC gene embeddings (Y-axis, mean cosine similarity) versus a traditional co-expression network (X-axis, mean Pearson correlation) within known biological pathways (KEGG). Each point represents a KEGG pathway. All points lie above the y=x diagonal (red dashed line), demonstrating that scDMC more accurately captures functional associations between genes. **c, d**, Functional enrichment analysis of the gene modules identified in CD8+ T cells (c) and HSCs (d). The size of the points represents the proportion of genes in the module involved in the pathway, while the color intensity indicates the statistical significance of the enrichment (-Log10(P-value)). The results show that each module corresponds to a specific biological function. **e**, Cell-type-specific gene regulatory networks inferred from scDMC embeddings. The left panel shows the sub-network centered on the key metabolic gene SPX3 in CD8+ T cells. The right panel shows the sub-network centered on CCDC144A in HSCs, whose neighbors are enriched in key factors for maintaining stem cell function. **f**, Heatmap of attention weights from scDMC during cell type annotation on the PBMC10k dataset. Each row represents a high-attention gene, and each column represents a randomly sampled cell, colored by the cell type annotation at the top. The heatmap clearly shows that when classifying different cells, the model’s attention focuses on corresponding, biologically meaningful marker genes. **g, h**, Ablation study of scDMC’s key components. The line plot in g shows the progressive performance improvement on cell type annotation (F1-Score) and batch integration (AvgBio score) tasks on two representative datasets (Myeloid and Immune human) as model complexity increases (from the rank-only RankOnly to the full scDMC). The bar plot in h more clearly quantifies the performance gain brought by each component for the F1 and AvgBio metrics. Together, the results demonstrate that the introduction of the dual-stream mechanism and momentum contrastive learning provides a critical, synergistic benefit to the model’s final performance.

To further and more objectively assess the quality of the gene modules discovered by scDMC, we selected 235 pathways with more than 10 genes from the KEGG database as a “gold standard” gene set. For each pathway, we calculated the average cosine similarity of all gene pairs within the pathway in the scDMC embedding space, as well as their average Pearson correlation coefficient in the gene co-expression matrix. We demonstrate that the gene embeddings learned by scDMC are significantly superior to traditional Pearson correlation-based co-expression analysis in capturing functional associations between genes. When evaluated for consistency within known KEGG biological pathways, scDMC exhibited higher internal similarity across all tested pathways (Fig. 5b), indicating that scDMC can more accurately and robustly reflect true functional gene relationships.

Leveraging these high-quality gene embeddings, we constructed cell-type-specific gene regulatory networks (GRNs) and identified key regulatory factors (hub genes) within them. For example, in CD8+ T cells, we identified the critical metabolic gene SPX3 (SLC37A3) as a core hub gene. In HSCs, we identified the functionally uncharacterized CCDC144A as another core hub. Analysis of the sub-networks formed by these hub genes and their neighbors revealed their central roles in coordinating critical cellular functions, such as T-cell metabolic reprogramming and HSC stemness maintenance (Fig. 5e).

### 2.6 scDMC’s decision-making is biologically interpretable

To open the “black box” of the model, we conducted an in-depth exploration of scDMC’s internal working mechanisms during downstream tasks. The results show that scDMC’s decision-making process is not only interpretable but also highly consistent with established biological knowledge.

The core of a Transformer model is its attention mechanism. In the cell type annotation task, the attention weight assigned to a specific gene for correctly classifying a cell can reflect that gene’s importance for identifying that cell type. We extracted the attention scores for all genes in each cell and sampled them by cell type to identify high-attention genes for each class.

By analyzing the attention weights assigned to genes during the classification task, we found that scDMC exhibits a “thinking process” akin to that of a domain expert. In the PBMC10k dataset, high-attention genes showed a strong overlap with core functional markers for specific cell types (Fig. 5f). For example, the “cytotoxicity toolkit” of CD8+ T cells (e.g., NKG7, PFP) and the key transcription factor for B cells, SPIB, were all assigned extremely high attention. This indicates that scDMC makes accurate classifications by focusing on the most biologically relevant genes.

More interestingly, we discovered that the fine-tuning process not only optimizes attention weights but also reshapes the gene embedding space to reflect the functional programs of specific cell types. Through unsupervised clustering of the fine-tuned gene embeddings, we identified multiple “gene programs” whose activation patterns showed high specificity across different cell types (Supplementary Fig. 3). For instance, the synergistic activation of numerous gene programs observed in dendritic cells perfectly illustrates their “multi-threaded” working mode as “professional” antigen-presenting cells. These findings demonstrate that scDMC is not only superior in performance but also that its internal mechanisms have a solid, explorable biological foundation.

### 2.7 Ablation study reveals the synergistic contribution of scDMC components

To systematically dissect the contributions of each core component within the scDMC framework, we conducted a series of ablation studies. We progressively added key components: starting with a rank-only encoding (RankOnly), then adding expression-aware encoding (EPEOnly), enabling the dual-stream (Dual), and finally the full scDMC to observe the performance improvements. All variant models were pre-trained using the exact same 2 million single-cell dataset and training strategy as the full scDMC.

The experimental results clearly demonstrate the synergistic effect and indispensability of each component (Fig. 5g, 5h). The performance improvement from the single-stream RankOnly to EPEOnly validates the superiority of our proposed expression-aware encoding strategy. The subsequent introduction of the dual-stream mechanism (Dual) further enhanced performance, indicating that simultaneously modeling the relative importance and absolute magnitude of gene expression allows the model to learn more comprehensive cellular features.

Most critically, the complete scDMC model (i.e., adding momentum contrastive learning on top of the dual-stream) achieved the best performance across all evaluation metrics, with the most significant margin of improvement. This decisively proves that the introduction of cell-level momentum contrastive learning plays a crucial synergistic role. The dual-stream architecture provides natural, high-quality positive pairs for contrastive learning, which in turn drives the model to learn more robust and discriminative global cell representations.

## 3 Discussion

Single-cell sequencing technologies are leading a revolution in biomedicine, yet extracting deep biological principles from the massive, high-dimensional, and sparse data they generate remains a formidable computational challenge. Foundation models offer a highly promising solution; however, existing single-cell language models still face fundamental bottlenecks in information fidelity and the comprehensiveness of representation learning. In this work, we have introduced scDMC, a next-generation single-cell foundation model designed to directly address these core challenges through its innovative dual-stream pre-training framework.

Our research systematically demonstrates scDMC’s superior performance and profound biological insight. At the cellular level, it sets a new state-of-the-art (SOTA) in multiple key tasks, including cell type annotation, unsupervised clustering, and batch effect integration. The success of scDMC is rooted in its unique architectural design. The dual-stream masked language modeling (Dual-Stream MLM) mechanism, by processing ranked and expression-aware information in parallel, captures both the relative functional context and the absolute expression magnitude of genes within a single, unified model for the first time, fundamentally resolving the information loss caused by single-encoding schemes. Furthermore, cell momentum contrastive learning leverages the natural positive pairs generated by the dual streams to drive the model toward learning a more discriminative global cell representation, which is the key to its exceptional robustness when handling rare cell types and complex datasets.

More importantly, scDMC is not merely a high-performance “black box”. We have delved deeply into the embedding space it learns, proving that it holds profound and interpretable biological meaning. It can deconstruct complex cellular functions into highly cohesive functional gene modules in a completely data-driven manner and further infer cell-type-specific regulatory networks centered around key hub genes. These discoveries not only align closely with established biological knowledge but also have the potential to uncover novel regulatory mechanisms. For instance, identifying a critical metabolic gene as the central hub of a CD8+ T cell regulatory network provides new computational evidence for the intimate link between immune cell function and metabolic reprogramming. This ability to distill interpretable biological hypotheses from massive datasets is the core value of next-generation foundation models in driving scientific discovery.

Despite its significant success, our work has limitations that also point to future research directions. First, the current scDMC is primarily based on transcriptomic data, whereas cellular function is determined by molecules at multiple levels. Our framework is designed with high extensibility; in the future, new input streams can be designed to integrate multi-modal data (such as protein information from CITE-seq or chromatin accessibility from ATAC-seq) to build a more comprehensive cell representation model. Second, we pre-trained our model on only 2 million human blood cells, which has already demonstrated the high efficiency of our framework. In the future, applying scDMC to larger-scale, cross-tissue, and even cross-species datasets for pre-training could enable it to learn more universal “rules of life”. Finally, applying the high-quality representations learned by scDMC to more complex downstream tasks, such as predicting drug responses, inferring cell differentiation trajectories, and modeling disease progression will be a key avenue for realizing its greater translational medicine value.

## 4 Methods

### 4.1 Data Processing and Dual-Stream Input Construction

Single-cell sequencing data, after collection and organization, form a matrix with cells as rows and genes as columns, denoted as **X** ∈ ℝ^*N ×G*^. In single-cell RNA sequencing (scRNA-seq) data, each element **X**_*ij*_ ∈ ℝ^+^ represents the read count of an RNA molecule. This element **X**_*ij*_ denotes the RNA abundance of gene *j*∈ { 0, 1, …, *G*} in cell *i* ∈ {0, 1, …, *N*}. Hereafter, we refer to this matrix as the raw count matrix **X**.

**Vocabulary Construction.** In the scDMC framework, genes are treated as the fundamental units, or “tokens”, of the language model. We constructed a shared vocabulary containing 60,534 tokens, comprising 60,530 unique genes identified by their Ensembl IDs and four special tokens: **[PAD]** for padding, **[MASK]** for masking, **[CLS]** for global cell representation, and **[EOS]** to mark the end of a sequence. By uniformly mapping gene names from different datasets to their Ensembl IDs, we created a shared vocabulary that accommodates the union of all genes under study, thereby enabling the effective integration of heterogeneous data.

### 4.2 Dual-Stream Input Sequence Generation

To extract richer information from a single cellular expression profile, we generate two parallel input sequences for each cell. The constructed dataset is stored in the efficient Apache Parquet format, containing fields such as **S**_rank_, **S**_epe_, and **S**_value_, which are directly used for model pre-training.

**Input Stream 1: Ranked Gene Sequence (input ids rank).** This input stream is designed to capture the relative importance of gene expression within a cell, i.e., the contextual relationships between genes. We adopted and optimized the rank-based encoding method from Geneformer, with the following procedure:

- **Intra-cell Normalization (CP10k):** To eliminate the influence of sequencing depth, we normalize the gene counts for each cell to “Counts Per 10, 000”, yielding the matrix **X**^*′*^.

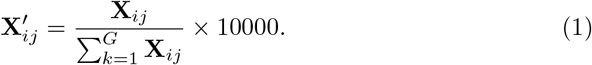
- **Gene Baseline Normalization:** To account for the inherent expression abundance of genes, we normalize **X**^*′*^ using a pre-computed median of non-zero expression for each gene, **Y**_*j*_, resulting in the matrix **X**^*′′*^. The value 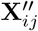 reflects the expression of gene *j* in cell *i* relative to its “typical” expression level across a large-scale single-cell dataset.

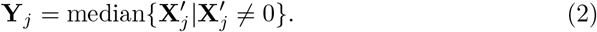

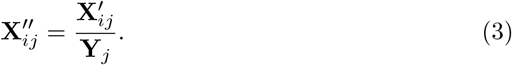
- **Ranking and Sequence Construction:** For each cell, we rank all genes with non-zero expression in descending order based on their values in **X**^*′′*^. The sequence of ranked gene IDs is then combined with special tokens to form the final input sequence **S**_rank_, which is padded or truncated to a pre-defined maximum length *M*.

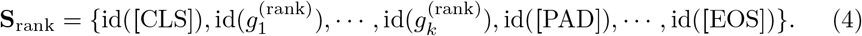

where 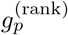 is the gene at the *p*th position after ranking, and *k* is the length of the non-zero expression sequence for the current cell *i*. This rank-based encoding method inherently normalizes the data, mitigating the effects of technical noise and batch effects. It also better captures the relative biological importance of genes in a specific cellular state, thereby helping the model learn regulatory relationships and deep cellular representations.

**Input Stream 2: Expression-Aware Sequence (input ids epe).** The expression-aware sequence serves as another input stream for the same cell, designed to preserve the absolute magnitude of gene expression and to be complementary to the first stream.

This sequence contains the exact same gene IDs as the first stream but maintains a fixed, unranked, original gene order (based on vocabulary index). Similarly, we retain only genes with non-zero expression and pad or truncate the sequence to the maximum length *M*, yielding the sequence Sepe **S**_epe_.

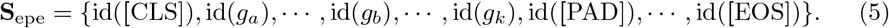

where *g*_*a*_, *g*_*b*_, … are the non-zero expressed genes appearing in vocabulary order, and *g*_*k*_ is the last non-zero gene in vocabulary order within cell *i*.

A key aspect of the expression-aware enhanced pre-training task is using the processed expression values to augment the model’s perceptual learning of the single-cell sequence data. Therefore, after constructing **S**_epe_, the gene expression values must be extracted and transformed. In parallel with the **S**_epe_ sequence, we prepare a corresponding expression value weight for each of its gene tokens. This weight is derived from the normalized matrix **X**^*′*^ (Eq. 1) and is transformed using a log 1*p* function to smooth the data distribution, resulting in the expression value matrix **V**:

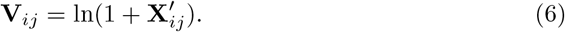

For each gene in **S**_epe_, we extract its corresponding **V**_*ij*_ value to form an expression value sequence, **S**_value_, of the same length as **S**_epe_. The weight for special tokens is set to 1.0. This **S**_value_ will be used as a weight in the subsequent MLM loss calculation.

The expression-aware sequence retains information about the “level of gene expression”, a representation that highlights the magnitude and absolute importance of genes within a single cell. This emphasizes “which genes are more important”, thereby directing the model’s focus during learning and providing it with a more comprehensive learning perspective.

### 4.3 scDMC Model Architecture

The scDMC model adopts the Transformer as its base architecture, as illustrated in Fig. 1, consisting of two weight-sharing online encoders and one momentum encoder. The online encoders are responsible for processing the two aforementioned input streams, while the parameters of the momentum encoder are an exponential moving average (EMA) of the online encoder’s parameters. The momentum encoder serves to provide a more stable and consistent target representation for the contrastive learning task.

### 4.4 Pre-training Objectives

The design of multiple pre-training objectives is one of the main innovations of scDMC. These objectives are crafted to enable the model to capture the contextual information of gene regulatory networks while simultaneously learning a representation for the entire single-cell data sequence, thereby better discriminating subtle differences between cells. The pre-training of scDMC is driven by two core self-supervised tasks: Dual-Stream Masked Language Modeling (Dual-Stream MLM) and Cell Momentum Contrastive Learning. These two tasks work synergistically to allow the model to learn comprehensive and robust biological representations at both the gene level and the whole-cell level.

**Dual-Stream Masked Language Modeling.** The dual-stream MLM task enables the model to learn contextual dependencies and co-expression patterns among genes. The rank-based encoding more accurately captures the relative biological importance of genes in a specific cellular state, while the expression-aware encoding highlights the magnitude and absolute importance of gene expression within a single cell. These two streams are complementary and indispensable, helping the model to learn regulatory relationships between genes and deep cellular representations.

We perform masked language modeling independently on the two input streams, **S**_rank_ and **S**_epe_. For each input sequence, we randomly select 15% of the non-special tokens to mask. This dual-stream masking strategy serves two key purposes: (1) it forces the model to learn robust cross-feature relationships within each stream, preventing over-reliance on either stream alone, and (2) the independent corruption of each stream ensures the model can recover missing information by leveraging complementary biological signals from the unmasked portions. Following the strategy of BERT, 80% of the selected tokens are replaced with the **[MASK]** token, 10% are replaced with a random token from the vocabulary, and the remaining 10% are left unchanged. The model’s objective is to predict the original identity of these masked tokens based on the unmasked tokens in the sequence.

For the ranked gene sequence **S**_rank_, we calculate the standard cross-entropy loss. Let *θ* represent the parameters of the online encoder, 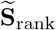 be the masked sequence, and *m* be the set of indices for the masked tokens. The ranked-stream MLM loss, 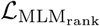, is thus defined as:

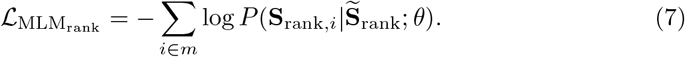

This ranked-stream MLM loss encourages the model to learn the relative importance ranking and interrelationships of genes within the cell.

For the expression-aware sequence **S**_epe_, we introduce a key innovation: an expression-weighted loss, 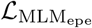. We utilize the weight sequence **S**_value_, which is extracted from the log 1*p* transformed expression value matrix **V**, to adjust the contribution of each masked token to the loss calculation. Specifically, the weighted loss is defined as:

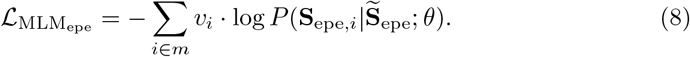

where *v*_*i*_ is the expression value weight corresponding to the *i*th masked token in the sequence. This weighting scheme ensures that the model is penalized more heavily for incorrectly predicting highly expressed genes. As a result, the model is encouraged to not only focus on gene co-occurrence relationships but also to recognize the absolute magnitude of gene expression. This approach allows the model to learn more refined information about the cellular state.

**Cell Momentum Contrastive Learning.** To enhance the model’s ability to discriminate at the cell (“sentence”) level, we introduce cell momentum contrastive learning. The goal of this task is to learn a high-dimensional embedding space where representations from different views of the same cell (i.e., the two input streams) are pulled closer together, while representations from different cells are pushed further apart. Specifically, we use the final hidden state at the position of the **[CLS]** token from each sequence as the global representation of the cell for contrastive learning. This design encourages the model to develop a more effective pooling mechanism directly within the Transformer blocks, leading to more robust and discriminative cell representations.

Let **q**_rank_ and **q**_epe_ be the representation vectors corresponding to the **[CLS]** special token, obtained after the online encoder (with parameters *θ*_*q*_) has processed the masked sequences 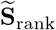 and 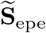, respectively. Concurrently, let **k**_rank_ and **k**_epe_ be the **[CLS]** representation vectors obtained after the momentum encoder (with parameters *θ*_*k*_) has processed the same input sequences. We construct the contrastive loss in two directions. The first direction treats **q**_rank_ as the query and **k**_epe_ as the positive key. The second direction treats **q**_epe_ as the query and **k**_rank_ as the positive key. The **[CLS]** representations output by the momentum encoder for other cells within the same batch constitute the negative keys. The contrastive loss takes the form of InfoNCE [54], and for a single positive pair (*q, k*^+^), the loss is calculated as follows:

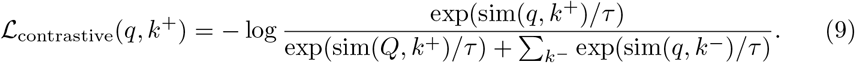

where 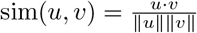 is the cosine similarity, and *τ* is a temperature hyperparameter that controls the smoothness of the distribution.

The total contrastive loss for scDMC ℒ_CellMoCo_, is the average of the losses from both directions:

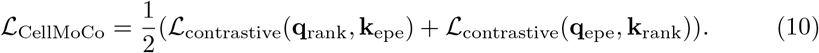

The parameters of the momentum encoder, *θ*_*k*_, are not updated via backpropagation. Instead, they are updated as an exponential moving average (EMA) of the online encoder’s parameters, *θ*_*q*_:

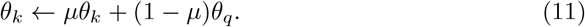

where *µ* ∈ [0, 1) is the momentum coefficient, set to 0.995 in our experiments. This “momentum” update mechanism ensures the smoothness and consistency of the cell representations.

**Overall Loss Function.**The pre-training objective of scDMC combines the masked language modeling (MLM) and cell momentum contrastive learning (Cell-MoCo) losses through a weighted summation:

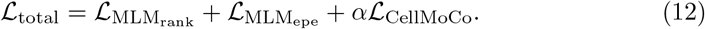

where *α* is a task-balancing hyperparameter that controls the relative contribution of the contrastive learning loss ( ℒ_CellMoCo_) to the overall objective. In our experiments, we set *α* = 1 to assign equal importance to both the MLM and contrastive learning tasks, ensuring neither objective dominates during optimization. By jointly optimizing this multi-task objective, scDMC can simultaneously capture rich biological information at both the gene and cell levels, thereby learning more refined cellular information and high-quality cell representations.

### 4.5 Efficient Training Strategies

The quadratic computational complexity of the standard Transformer architecture is a primary bottleneck when processing the long gene sequences common in single-cell data, which can have a maximum length of 2,048 in scDMC. To ensure the computational feasibility and training efficiency of our framework, we integrated three key architectural optimizations.

**Efficient Attention Mechanism.** We replaced the standard self-attention mechanism with FlashAttention. By optimizing the I/O (Input/Output) of the GPU memory, FlashAttention significantly reduces memory usage and accelerates the computation process without any approximation. This is crucial for handling long gene sequences with limited computational resources.

**Dynamic Sequence Length Processing.** To further enhance training efficiency, our custom data collator is capable of handling sequences of dynamic length. It generates an attention mask for each batch, which ensures that the model ignores the padded **[PAD]** tokens during the attention calculation. This strategy avoids redundant computation on the non-informative parts of the sequence, thereby increasing the overall training throughput.

**Positional Information via ALiBi.** Given that the order of genes in a single-cell expression profile has no inherent linear biological meaning, traditional absolute or relative positional encodings are not an optimal choice. We therefore adopted Attention with Linear Biases (ALiBi) to encode positional information. We posit that ALiBi can more effectively capture the non-sequential, complex regulatory relationships between genes and can better generalize to input sequences of varying lengths.

The core mechanism of ALiBi is to discard traditional learnable positional embedding vectors. Instead, it adds a static, non-learnable bias term directly to the attention scores (the query-key dot product) *after* they are calculated but *before* the softmax function is applied. The magnitude of this bias term is linearly proportional to the relative distance between the query and key tokens. Specifically, for each attention head, a pre-set scalar slope*m* is used, and the bias value applied to the attention score is calculated as *m* · |*i* − *j*|, where *i* and *j* are the positions of the query and key tokens, respectively. This method effectively injects a positional inductive bias directly into the attention mechanism. Finally, after the ALiBi adjustment and normalization with the softmax function, we obtain the attention weights *v*^*^.

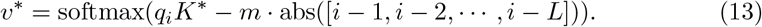

## Data availability

All data used in this study are publicly available. Specific usage details and data descriptions can be found in Supplementary Information S5. The pretraining dataset was obtained from the CELLxGENE Census database, release version dated 2024-07-01 (https://cellxgene.cziscience.com/). The Zheng68K dataset was downloaded from the “Fresh 68k PBMCs” section of the 10x Genomics Single Cell Gene Expression database (https://www.10xgenomics.com/ datasets/fresh-68k-pbmcs-donor-a-1-standard-1-1-0). The PBMC10K dataset was retrieved from the scVI-tools package (https://scvi-tools.org/) using the scvi.data.pbmc dataset API. The BMMC dataset is available from the Gene Expression Omnibus (GEO) database under accession number GSE194122 (https://www.ncbi.nlm.nih.gov/geo/query/acc.cgi?acc=GSE194122). The Immune Human dataset was obtained from Figshare (https://figshare.com/articles/dataset/ Benchmarking atlas-level data integration in single-cell genomics - integration task datasets Immune and pancreas /12420968/8). The MS dataset was obtained from https://www.ebi.ac.uk/gxa/sc/experiments/E-HCAD-35/downloads. The Myeloid dataset is publicly available from the GEO database under accession number GSE154763.

## Code availability

The source code, pre-trained models, and datasets for this study have been made publicly available to ensure reproducibility and facilitate further research: Implementation Code: https://github.com/HonglieGuo/scDMC. Datasets: https://huggingface.co/datasets/Honglie/scDMC-datasets. Models: https://huggingface.co/Honglie/scDMC-models/tree/main

## Acknowledgements

We thank the anonymous reviewers for their valuable suggestions. This work was supported by the National Natural Science Foundation of China under Grants 62462068, 62062067.

## Author contributions

Honglie Guo, Qinghang Cui and Xiang Zhang conceived the study. Honglie Guo, Qinghang Cui and Chaowei Chen collected the downstream datasets involved in this article. Honglie Guo, Qinghang Cui developed data collection criteria and strategies for pre-training. Honglie Guo, Qinghang Cui and and Shunfang Wang proposed the pretraining framework. Honglie Guo, Qinghang Cui and Xiang Zhang implemented and pre-trained the models. Honglie Guo, Qinghang Cui and Xiang Zhang benchmarked all methods. Honglie Guo, Qinghang Cui, Weihua Zheng and Changfeng Cai provided a lot of advice on pre-training framework design and downstream tasks. Honglie Guo, Qinghang Cui and Xinyi Wang wrote the manuscript. All authors read and approved the final manuscript.

## Additional information

Evaluation Metrics & Supplementary Figures & Supplementary Tables

