## Supplementary materials for scDMC for "Unlocking biological insight from single-cell data with an interpretable dual-stream foundation model" (from another version of this preprint: version 1, 2025-09-11)

#### Supplementary Notes

#### S1. Model Settings and Pre-training Process

The scDMC model consists of 12 Transformer layers, each with 8 attention heads, and a hidden embedding size of 512. The extrapolation size for the ALiBi linear bias attention was set to 2,048, and the vocabulary size was 60,534. During training, a hidden dropout probability of 0.02 and an attention dropout probability of 0.1 were used to prevent overfitting. The model was implemented in PyTorch, utilizing the Hugging Face Transformers library for model configuration, data loading, and training.

The pre-training process for scDMC was conducted on a single A800 GPU, taking approximately 10 days to train for 3 epochs on the 2 million single-cell dataset. We used the AdamW optimizer with a GeLU activation function, an initial learning rate of 5e-5, and a batch size of 24. Upon completion of training, the model achieved a best loss value of 2.93 on the validation set. The final model parameters were saved to a specified checkpoint path, ready for subsequent fine-tuning or inference on downstream tasks.

The pre-training process shows the evolution of four key loss components of the scDMC model. The subplots correspond to the change over training steps for (top left) MLM_rank Loss, (top right) MLM_epe Loss, (bottom left) Contrastive Loss, and (bottom right) Total Loss. All loss components exhibit a healthy downward trend, indicating that the model is learning effectively and converging. To clearly observe the dynamics of the MLM_epe Loss, whose magnitude is much smaller than the other losses, a logarithmic scale is used for its Y-axis. The stable decline of all metrics confirms the stability and effectiveness of the pre-training strategy employed by the scDMC model.

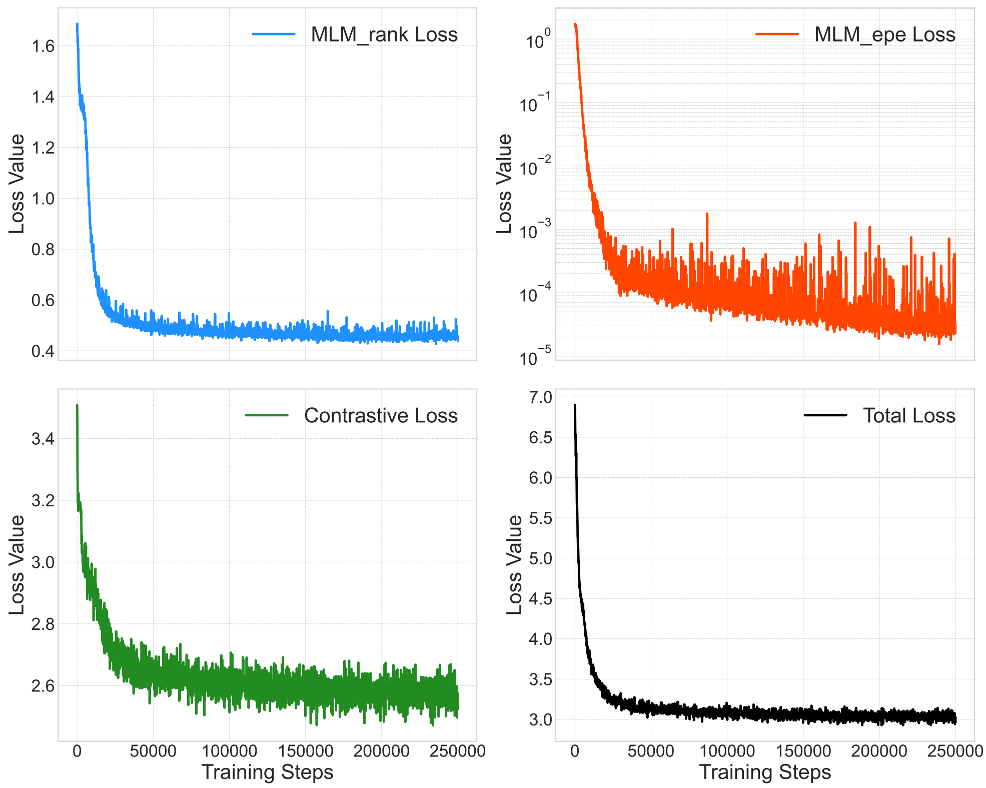


**Supplementary Fig. 1**: Analysis of loss components during the scDMC pre-training process.

### S2. Detailed workflow for gene module discovery and functional enrichment analysis

To identify functionally related gene modules, we employed the following workflow, which was applied to two representative cell types:

**(1) Gene Embedding Extraction.** First, from an scDMC model fine-tuned on a specific dataset (e.g., Zheng68K), we extracted the average contextual gene embeddings for all expressed genes (12,535 in total) within a specific cell type (e.g., CD8+ cytotoxic T cells). These embedding vectors were organized into a feature matrix with genes as rows and embedding dimensions as columns.

**(2) Gene Module Clustering.** Subsequently, we applied the unsupervised graph-based clustering algorithm Leiden (from the leidenalg Python package, with the resolution parameter set to 0.2) to the aforementioned gene embedding matrix. This was done to identify functionally related groups of genes, or “gene modules”. In CD8+ T cells, this procedure identified 8 major gene modules (Main Fig. 5a).

**(3) Functional Enrichment Analysis.** We used the online gene enrichment analysis tool Metascape (https://metascape.org/) to functionally annotate each identified gene module. For the largest module identified in CD8+ T cells (Module 0, containing 3,836 genes), its size exceeded Metascape's input limit of 3,000 genes. We inferred that this module likely represents housekeeping genes involved in basic cellular activities with dispersed functions. Therefore, it was excluded from functional enrichment analysis, and we focused on the remaining 7 more functionally specific modules (Main Fig. 5c).

**(4) Cross-Dataset Validation.** To validate the robustness of this workflow, we repeated steps 1 through 3 verbatim on hematopoietic stem cells (HSCs) from the BMMC dataset. The results similarly identified gene modules with high functional cohesion (Main Fig. 5d), demonstrating the generalizability of our analytical approach.

### S3. Gene Regulatory Network (GRN) Construction and Hub Gene Identification

A central challenge in constructing a GRN is determining an appropriate similarity threshold. To address this, we systematically evaluated how key network topological properties evolve as a function of the cosine similarity threshold, ranging from 0.5 to 0.9 (Supplementary Fig. 2). The objective of this analysis was to identify an optimal threshold that effectively filters out weak connections (likely noise) while preserving the core network of strong, biologically meaningful interactions.

In two distinct cell types and datasets, we consistently observed a clear "phase transition" region with a critical point near the similarity threshold of 0.7 (marked by a red dashed line) (Supplementary Fig. 2). At this critical point, the number of edges (Top) and the degree of modularity (Middle) both decrease precipitously, indicating the removal of a vast number of weak-to-moderate connections. Crucially, while the largest connected component (Bottom) also contracts sharply, it stabilizes into a substantial core network rather than completely disintegrating. This phenomenon indicates that the 0.7 threshold acts as a critical inflection point for separating strong biological signals from noise. Therefore, based on this robust, cross-dataset observation, we uniformly adopted 0.7 as the similarity threshold for all GRN analyses in the main text. Hub genes within these networks were subsequently identified by calculating their degree centrality (Supplementary Table 1).

### S4. Discovering and Validating Functional Gene Programs from Fine-Tuned Gene Embeddings

To validate our core hypothesis that the fine-tuning process refines the model's understanding of gene semantics, we designed an analysis to discover functionally related "gene programs" from fine-tuned gene embeddings in a data-driven manner. Specifically, from an scDMC model that was fine-tuned for a particular downstream task (e.g., cell type annotation), we extracted the contextual embeddings for all genes. Subsequently, we applied the unsupervised clustering algorithm Leiden to group these

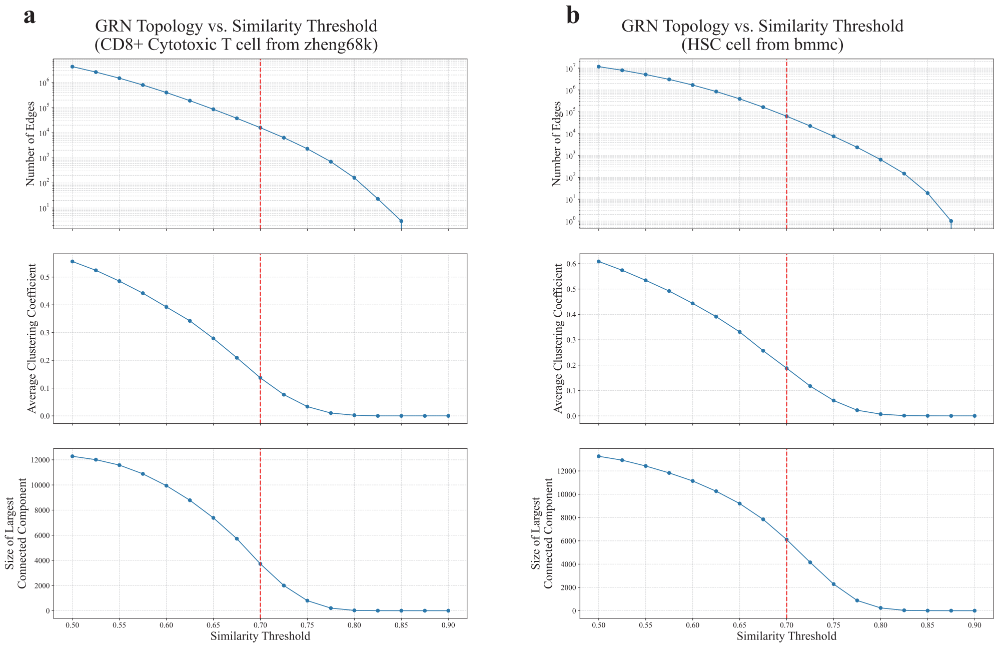


**Supplementary Fig. 2**: Selecting the similarity threshold for Gene Regulatory Networks (GRNs) based on network topology. **a**, Analysis performed on the GRN from CD8+ cytotoxic T cells in the Zheng68k dataset. **b**, The same analysis performed on the GRN from hematopoietic stem cells (HSCs) in the BMMC dataset.In each analysis panel, the plots from top to bottom show: **Number of Edges:** Plotted on a logarithmic scale, reflecting network density. **Average Clustering Coefficient:** A measure of network modularity or the tendency of nodes to form tightly connected groups. **Size of Largest Connected Component:** Reflects the main connectivity of the network.

gene embeddings. To evaluate the biological significance of these data-driven gene programs, we calculated the average activation score (Z-score) for each program across various cell types and visualized the results as a heatmap (Supplementary Fig. 3).

The analysis revealed highly specific associations between gene programs and cell types, which are in strong agreement with established biological knowledge. In the Zheng68k dataset (Supplementary Fig. 3a), the synergistic, high activation of numerous gene programs in Dendritic Cells perfectly reflects their "multi-threaded" working mode as "professional" antigen-presenting cells. Furthermore, the specific activation of Program 16, which includes the WNT16 gene, in CD34+ cells demonstrates the model's ability to capture core regulatory pathways essential for maintaining stem cell identity.

This finding was validated in the independent PBMC10k dataset (Supplementary Fig. 3b). A particularly striking example is the strong activation of nearly all gene

**Supplementary Table 1:** List of the Top 10 Hub Genes in CD8+ T cells.

| Rank | Gene Symbol | Degree Centrality |
|---|---|---|
| 1 | SPX3 | 0.0537 |
| 2 | FAM153A | 0.0491 |
| 3 | C9ORF34 | 0.0375 |
| 4 | TFIIS.H | 0.0373 |
| 5 | TBS2 | 0.0373 |
| 6 | MCE | 0.0350 |
| 7 | DISPA | 0.0348 |
| 8 | IP6K | 0.0331 |
| 9 | TAM41 | 0.0280 |
| 10 | ZNF439 | 0.0276 |

programs in Megakaryocytes, which accurately portrays their global transcriptional state, geared for massive biosynthesis as "platelet factories".

These analyses demonstrate that clustering the fine-tuned gene embeddings from scDMC can reliably identify gene programs that reflect the core identity and functional state of cells, providing a powerful tool for interpretable biological discovery using deep learning models.

### S5. Datasets

#### S5.1 Pre-training Dataset

The pre-training dataset was sourced from the CELLxGENE Census database and comprises 2 million healthy human blood cells. We collected the data for our pre-training model from the CELLxGENE portal (https://cellxgene.cziscience.com/), using the Census API (https://chanzuckerberg.github.io/cellxgene-census/python-api.html) to query for the "blood" tissue from the CELLxGENE Census (release version 2024-07-01), downloading a total of over 5.4 million blood single-cell records. To demonstrate the efficiency of the scDMC framework under limited computational resources, we selected the first 2 million cells from this collection to construct our pre-training dataset using our tokenizer and vocabulary. We only standardized the gene expression matrix during the tokenization process, without performing any additional cell quality control or dimensionality reduction steps. The pre-training dataset includes a total of 131 different cell types from 6,603 healthy human donors, of whom 67.3% were male.

Detailed information is provided in Supplementary Fig. 4, showing that the primary cell populations are "CD14-positive monocyte", "natural killer cell", and "central memory CD4-positive, alpha-beta T cell". This clearly indicates that these three cell types are the most abundant and dominant populations in the sample. Secondary cell populations, such as "B cell" and "monocyte", have moderately sized boxes, indicating they also constitute a non-negligible proportion of the sample.

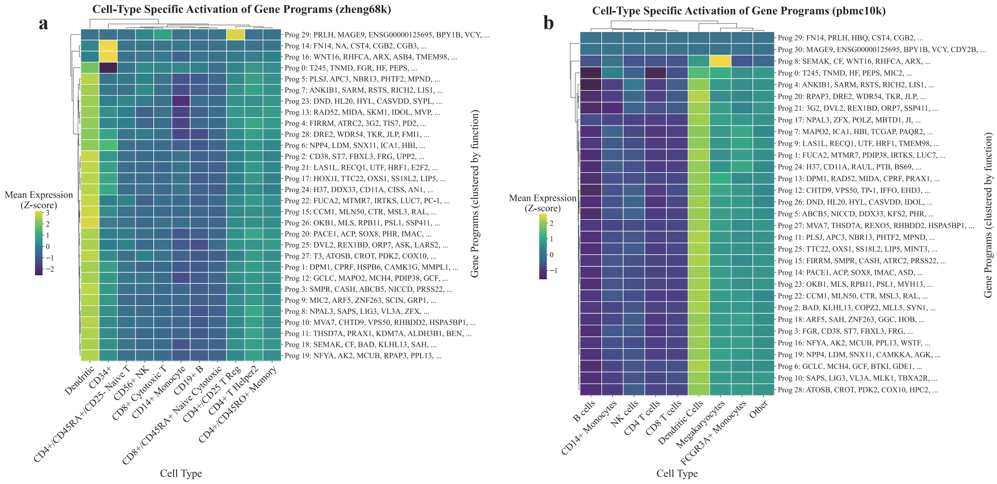


**Supplementary Fig. 3**: Cell-type-specific gene programs identified from fine-tuned scDMC embeddings. **a, b**, Heatmaps showing the average activation scores (Z-scores) of "gene programs" (rows), identified by unsupervised Leiden clustering on fine-tuned scDMC gene embeddings, across different cell types (columns) in two independent datasets: Zheng68k (**a**) and PBMC10k (**b**). Each row represents a data-driven gene program, and each column represents a cell type. Color intensity (yellow for high, purple for low) indicates the relative activation level of the gene program in that cell type. The heatmaps clearly show highly specific association patterns between gene programs and cell types. For example, in the Zheng68k dataset, numerous programs are synergistically activated in Dendritic Cells, and in the PBMC10k dataset, nearly all programs are activated in Megakaryocytes. These patterns are highly consistent with known cellular biological functions.

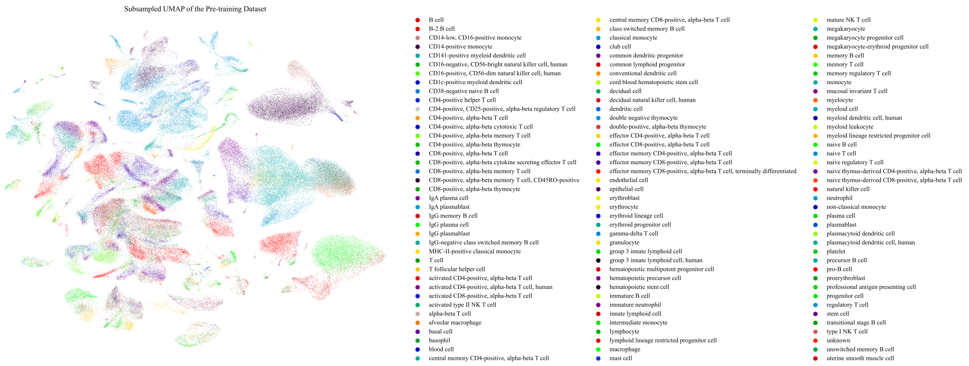


**Supplementary Fig. 4**: Visualization of cell heterogeneity in the pre-training dataset. The UMAP plot shows a 10% random subsample of cells, with different colors representing different cell type identities.

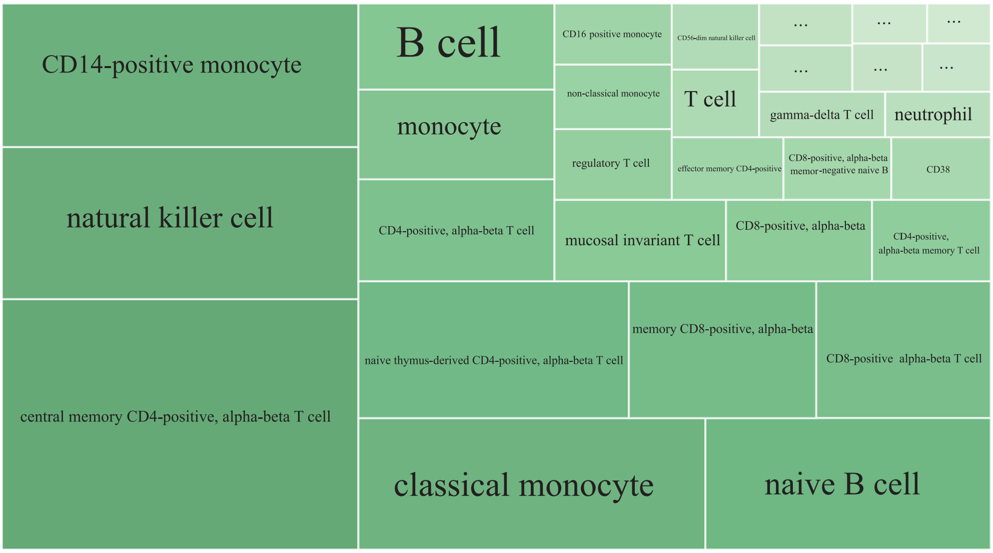


**Supplementary Fig. 5**: Distribution of major cell types in the pre-training dataset. Each rectangle in the treemap represents a specific cell type, with its area being proportional to the number or percentage of that cell type in the total sample.

#### S5.2 Downstream Task Datasets

- **Immune Human:** A comprehensive immune cell atlas containing 33,506 cells, integrating 5 scRNA-seq datasets from bone marrow and peripheral blood. This dataset encompasses multiple sequencing technologies (10x v2/v3, Smart-seq2), includes 16 cell types, and has 10 different technology batches, making it an ideal benchmark for evaluating model performance on cell type annotation and cross-platform batch integration.
- **Zheng68K:** A classic dataset of approximately 68,000 human peripheral blood mononuclear cells (PBMCs). It is renowned for its imbalanced cell type distribution and high similarity between certain subtypes, including several rare cell types (e.g., CD34+ progenitor cells), posing a stringent challenge to a model's rare-cell identification and fine-grained classification abilities.
- **PBMC10k:** A multi-batch dataset of approximately 12,000 PBMCs from two batches. It is commonly used to evaluate a model's ability to integrate batch effects. We test whether scDMC can maintain a clear distinction among 9 different immune cell subtypes while removing batch differences.
- **BMMC (CITE-seq):** A multimodal dataset of over 90,000 human Bone Marrow Mononuclear Cells. We used only its gene expression (GEX) component. Sourced from 12 different healthy donors and containing 45 finely annotated immune cell subtypes, it serves as a gold standard for testing model performance on multi-donor batch integration and ultra-high-resolution cell annotation.
- **MS (Multiple Sclerosis):** A single-cell dataset from brain tissue of multiple sclerosis patients and healthy controls. This dataset involves both healthy and diseased

states and includes samples from different brain regions (e.g., prefrontal cortex), making it a valuable resource for testing model robustness and representation capabilities on disease-related datasets.

- **Myeloid (Pan-cancer):** A pan-cancer myeloid cell dataset containing samples from tumor and adjacent normal tissues across 9 different cancer types. Following the setup of the original study, we divided it into a reference set (6 cancer types) and a query set (3 cancer types) to evaluate the model's generalization and annotation capabilities in complex scenarios across different cancer types and tissue sources.

**Supplementary Table 2:** Detailed information for downstream task datasets.

| Dataset | Tissue | Disease | Number of Batches | Number of Cell Types | Number of cells |
|---|---|---|---|---|---|
| BMMC | Marrow | Normal | 12 | 45 | 90261 |
| Immune | Marrow and Blood | Normal | 10 | 16 | 33506 |
| MS | Brain | Multiple Sclerosis | 2 | 18 | 21312 |
| Myeloid | Marrow | Pan-cancer | 2 | 21 | 13178 |
| PBMC10K | Blood | Normal | 2 | 9 | 11989 |
| Zheng68K | Blood | Normal | 1 | 11 | 68263 |

### S6. Evaluation Metrics

To comprehensively evaluate the performance of our model, we employed a variety of established evaluation metrics tailored to different downstream tasks. These metrics are primarily used to quantify and assess the model's performance in supervised cell type annotation, unsupervised cell clustering, and batch integration.

#### S6.1 Cell Type Annotation Metrics

Cell type annotation is a supervised classification task where the goal is to assign a predefined type label to each cell. We use standard classification metrics: Accuracy, Precision, Recall, and Macro F1-Score to evaluate the cell type assignment performance. Assuming the set of true classes in a dataset is C, for any class $c \in C$, we define True Positives (TPc), False Positives (FPc), and False Negatives (FNc).

- **Precision:** For a class c, precision measures how many of the samples predicted as class c truly belong to that class.

$$\text{Precision}_c = \frac{TP_c}{TP_c + FP_c} \tag{1}$$

- **Recall:** For a class c, recall measures how many of all true samples belonging to that class were correctly predicted.

$$\text{Recall}_c = \frac{TP_c}{TP_c + FN_c} \tag{2}$$

- **F1-Score:** The F1-Score is the harmonic mean of precision and recall, providing a combined measure of their performance.

$$\text{F1}_c = 2 \cdot \frac{\text{Precision}_c \cdot \text{Recall}_c}{\text{Precision}_c + \text{Recall}_c} \tag{3}$$

- **Macro F1-Score::** To evaluate the model's performance equally across all classes, especially in datasets with class imbalance, we calculate the Macro F1-Score. It is the arithmetic mean of the F1-Scores for all classes.

$$\text{Macro F1} = \frac{1}{|C|} \sum_{c \in C} \text{F1}_c \tag{4}$$

#### S6.2 Unsupervised Clustering and Batch Integration Metrics

For unsupervised clustering and batch integration tasks, we evaluate the quality of the cell embeddings generated by the model. These metrics are largely based on the scIB benchmark framework and assess performance from two complementary dimensions: conservation of biological structure and correction of batch effects.

First are three metrics for the conservation of biological structure, which measure the extent to which the cell embeddings retain meaningful biological differences, such as distinguishing between different cell types.

- **Adjusted Rand Index (ARI):** ARI measures the similarity between two data clusterings. Here, we compare the algorithm-generated cluster labels $K$ with the true cell type labels C. The range of ARI is $[-1, 1]$, where 1 indicates a perfect match and 0 indicates the result is no better than random assignment.

$$\text{ARI} = \frac{\text{RI} - E[\text{RI}]}{\max(\text{RI}) - E[\text{RI}]} \tag{5}$$

  where RI is the Rand Index and $E[\text{RI}]$ is its expected value.
- **Normalized Mutual Information (NMI):** NMI measures the mutual dependence between two label assignments, independent of the actual label values. Its range is, where 1 indicates perfect agreement between the two label assignments. Let the true labels be Y and the clustering results be C.

$$\text{NMI}(Y, C) = \frac{2 \cdot I(Y; C)}{H(Y) + H(C)} \tag{6}$$

  where $I(Y; C)$ is the mutual information, and $H(Y)$ and$H(C)$ are the entropies of the label distributions.

- **Cell Type Average Silhouette Width (ASW):** The silhouette width measures how tightly grouped a sample is within its own cluster compared to other clusters. For a single sample, its silhouette coefficient $s$ is defined as:

$$s = \frac{b - a}{\max(a, b)} \tag{7}$$

where $a$ is the mean distance from the sample to all other points in the same cluster, and $b$ is the mean distance from the sample to all points in the next nearest cluster. We average the silhouette coefficients (based on true cell type labels) for all cells to get $\text{ASW}_{\text{cell-type}}$. To normalize its range to, we perform the following transformation:

$$\text{ASW}_{\text{cell}} = \frac{\text{ASW}_{\text{cell-type}} + 1}{2} \tag{8}$$

A higher $\text{ASW}_{\text{cell}}$ score indicates better separation of cell types in the embedding space.

Next are several evaluation metrics for batch effect correction, specifically designed to assess batch integration tasks by measuring the extent to which the model removes technical batch differences.

- **Batch Average Silhouette Width (ASW):** Similar to the cell type ASW, this metric calculates the average silhouette width based on batch labels, $\text{ASW}_{\text{batch-label}}$. However, its ideal value is -1 (indicating perfect mixing of cells from different batches). We transform it into a metric with a range of, where a higher score represents better batch mixing.

$$\text{ASW}_{\text{batch}} = 1 - |\text{ASW}_{\text{batch-label}}| \tag{9}$$

- **Graph Connectivity:** This metric assesses whether cells of each cell type maintain their connectivity in a k-nearest neighbor (k-NN) graph. For each cell type c, we calculate the size of its largest connected component (LCC) and divide it by the total number of cells of that type, Nc. The final score is the average over all cell types.

$$\text{GraphConn} = \frac{1}{|C|} \sum_{c \in C} \frac{|\text{LCC}(G_c^{kNN})|}{N_c} \tag{10}$$

A higher score indicates that the local neighborhood structure of cells is better preserved after integration.

To obtain an overall performance assessment, we aggregate the above metrics to produce a final composite score:

$$\text{Overall Score} = 0.6 \cdot \text{AVG}_{\text{Bio}} + 0.4 \cdot \text{AVG}_{\text{Batch}} \tag{11}$$

where AvgBio is the average of the biological conservation metrics (ARI, NMI, and $\text{ASW}_{\text{cell}}$), and $\text{AVG}_{\text{Batch}}$ is the average of the batch correction metrics (ASWbatch,

GraphConn). The Overall Score balances the two objectives of conserving biological structure and removing batch effects.

### S7. Additional Experimental Results

#### S7.1 Additional Results for Cell Type Annotation

The cell annotation performance of scDMC on the Zheng68k dataset highlights how its unique dual-stream pre-training framework not only learns the features of major cell populations but also enhances the discrimination of low-frequency signals (i.e., rare cell types) through contrastive learning. This is clearly evidenced in the confusion matrix comparison. For rare cell types such as CD34+ cells, scDMC achieved not only a recall of 0.91 but also a high precision of 0.93, significantly outperforming other models and signifying the higher reliability of its predictions. Furthermore, when handling easily confusable classes (e.g., CD4+/CD45RO+ Memory cells), scDMC also demonstrated the most balanced performance, avoiding the significant performance shortcomings seen in other models for this class. This comprehensive advantage ultimately overcomes the bias that models tend to develop on imbalanced data, ensuring the precise identification of all cell subpopulations. In clinical or biological research, rare cell types (such as cancer stem cells or specific immunoregulatory cells) often play a critical role. Therefore, this capability of scDMC is not merely a technical improvement but also signifies its immense potential for application in translational medicine research.

**Supplementary Table 3:** Performance comparison of various models on the cell type annotation task.

| **Models** | **Zheng68K** | | **PBMC10K** | | **BMMC** | | **MS** | |
|---|---|---|---|---|---|---|---|---|
| | Acc | F1 | Acc | F1 | Acc | F1 | Acc | F1 |
| scBert | 0.7982 | 0.6952 | 0.9554 | 0.8613 | 0.8240 | 0.5425 | 0.8365 | 0.7316 |
| scGPT | 0.8077 | 0.6706 | 0.9575 | 0.8615 | 0.8301 | 0.5699 | 0.8329 | 0.6356 |
| Geneformer | 0.8215 | 0.7034 | 0.9604 | 0.8228 | 0.8202 | 0.5128 | 0.7703 | 0.6645 |
| Geneformer-95M | 0.8456 | 0.7378 | **0.9733** | 0.9432 | 0.8543 | 0.6075 | 0.8787 | 0.7898 |
| scFoundation | 0.7316 | 0.6297 | 0.9716 | 0.9422 | 0.8375 | 0.5925 | 0.8278 | 0.8057 |
| LangCell | 0.8552 | 0.7459 | 0.9722 | 0.9502 | **0.8540** | 0.6171 | 0.8705 | 0.8004 |
| scDMC | **0.8660** | **0.7674** | 0.9708 | **0.9522** | 0.8511 | **0.6689** | **0.8792** | **0.8278** |

#### S7.2 Additional Results for Batch Integration

According to the experimental results in Supplementary Table 4, the scDMC model demonstrates comprehensive and significant superiority in cell clustering and batch integration tasks across several other single-cell datasets (BMMC, Immune, PBMC10K, MS, Myeloid). The model not only far surpasses the LangCell and Geneformer models on key clustering metrics that measure the conservation of biological

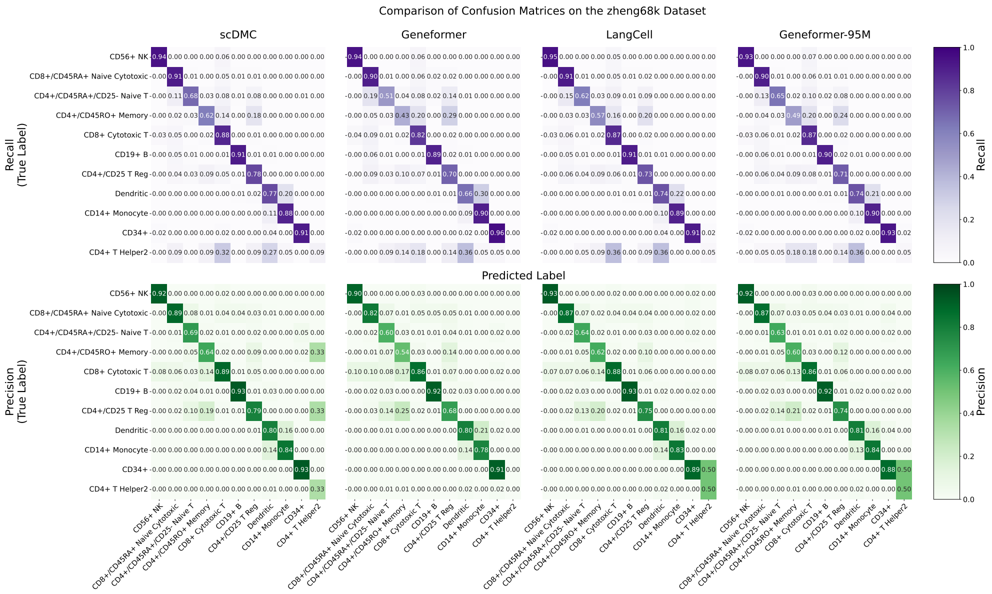


**Supplementary Fig. 6**: Performance of various models on the Zheng68k dataset, shown by normalized confusion matrices.

state, such as Adjusted Rand Index (ARI), Normalized Mutual Information (NMI), and cell type Average Silhouette Width ($ASW_{cell}$) achieving the most precise partitioning of cell subpopulations, but it also performs best on metrics that evaluate the removal of technical effects, such as the batch Average Silhouette Width ($ASW_{batch}$) and Graph Connectivity (GraphConn), effectively eliminating inter-batch differences. Ultimately, its Overall Score ranked first in all tests, fully demonstrating that scDMC can maximally preserve true biological heterogeneity while efficiently integrating multi-batch data.

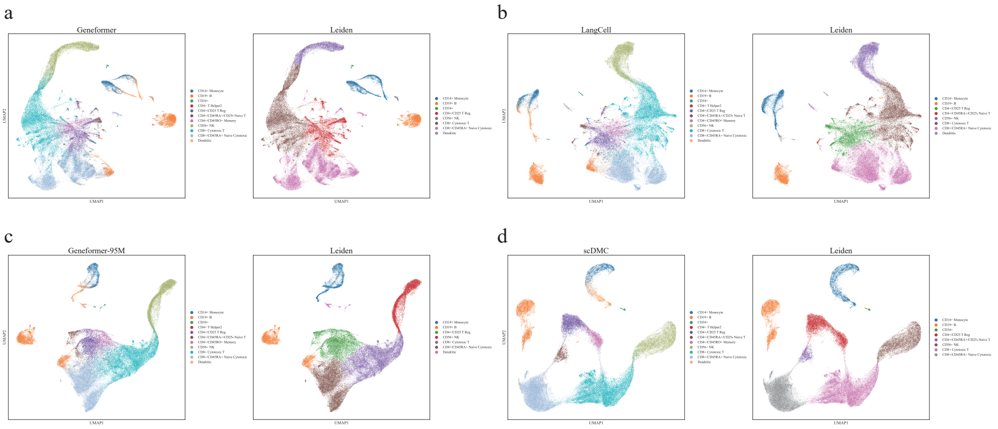


**Supplementary Fig. 7**: UMAP visualization of embedding clustering for different models on the Zheng68K dataset. The left panel is colored by model cell type embeddings (Ground Truth Labels). The right panel is colored by Leiden clustering results based on the embeddings (Leiden Clusters). **A**, Geneformer model. **B**, LangCell model. **C**, Geneformer-95M model. **D**, scDMC model.

**Supplementary Table 4:** Additional results for cell clustering and batch integration.

| **Datasets** | **Models** | **Metrics** | | | | | | | **Overall** |
|---|---|---|---|---|---|---|---|---|---|
| | | ARI | NMI | $ASW_{cell}$ | $Avg_{Bio}$ | $ASW_{batch}$ | GraphConn | $Avg_{batch}$ | |
| BMMC | Geneformer | 0.5786 | 0.7884 | 0.6632 | 0.6767 | 0.8774 | 0.9141 | 0.8957 | 0.7643 |
| | LangCell | 0.6037 | 0.8010 | 0.6812 | 0.6953 | 0.8886 | 0.9329 | 0.9107 | 0.7815 |
| | scDMC | **0.9548** | **0.9258** | **0.7959** | **0.8922** | **0.8994** | **0.9590** | **0.9292** | **0.9070** |
| Immune | Geneformer | 0.7681 | 0.7950 | 0.7473 | 0.7701 | 0.8602 | 0.9638 | 0.9120 | 0.8269 |
| | LangCell | 0.5340 | 0.7642 | 0.7612 | 0.6865 | 0.8695 | 0.9826 | 0.9261 | 0.7823 |
| | scDMC | **0.9331** | **0.9046** | **0.8201** | **0.8859** | **0.8700** | **0.9890** | **0.9295** | **0.9034** |
| PBMC10K | Geneformer | 0.9106 | 0.8490 | 0.8310 | 0.8635 | **0.9603** | 0.9440 | 0.9522 | 0.8990 |
| | LangCell | 0.8238 | 0.8706 | 0.8370 | 0.8438 | 0.9564 | 0.9848 | 0.9706 | 0.8945 |
| | scDMC | **0.9835** | **0.9689** | **0.9096** | **0.9540** | 0.9533 | **0.9907** | **0.9720** | **0.9612** |
| MS | Geneformer | 0.5027 | 0.6933 | 0.6285 | 0.6081 | 0.9229 | 0.9315 | 0.9272 | 0.7358 |
| | LangCell | 0.8191 | 0.8517 | 0.6997 | 0.7902 | 0.9258 | 0.9637 | 0.9448 | 0.8520 |
| | scDMC | **0.9440** | **0.9424** | **0.8404** | **0.9089** | **0.9422** | **0.9771** | **0.9596** | **0.9292** |
| Myeloid | Geneformer | 0.4511 | 0.5590 | 0.5576 | 0.5226 | 0.8799 | 0.8447 | 0.8623 | 0.6585 |
| | LangCell | 0.4783 | 0.6006 | 0.5780 | 0.5523 | 0.8954 | 0.8578 | 0.8766 | 0.6820 |
| | scDMC | **0.8970** | **0.8700** | **0.7769** | **0.8479** | **0.9113** | **0.9415** | **0.9264** | **0.8793** |

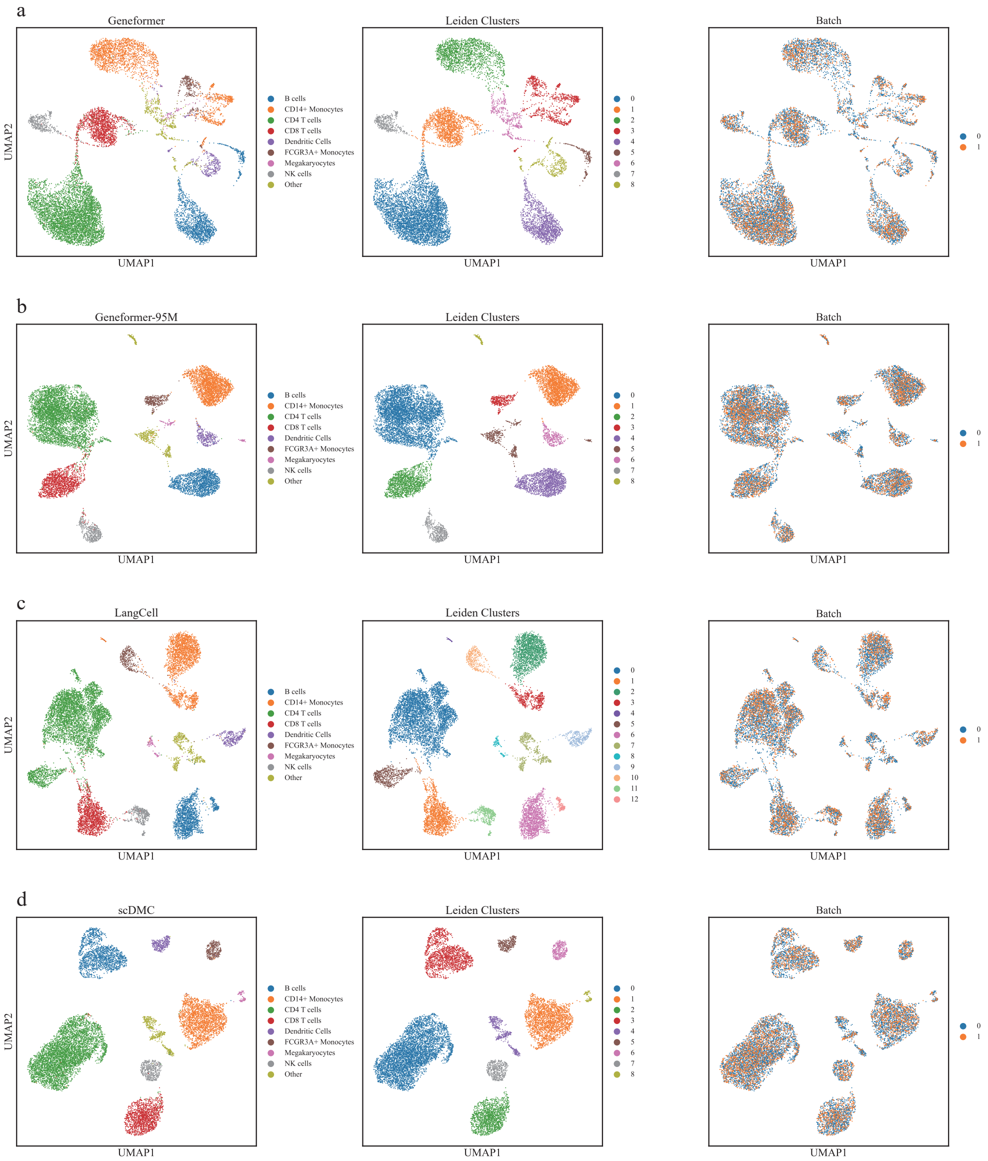


**Supplementary Fig. 8**: UMAP visualizations of cell embeddings for different models on the PBMC10K dataset. (Left) UMAP colored by Cell Type. (Middle) UMAP colored by Leiden clustering results from the model's embeddings (Leiden Clusters). (Right) UMAP colored by Batch. **a**, Geneformer. **b**, Geneformer-95M. **c**, LangCell. **d**, scDMC.
